# DetecDiv, a generalist deep-learning platform for automated cell division tracking and survival analysis

**DOI:** 10.1101/2021.10.05.463175

**Authors:** Théo Aspert, Didier Hentsch, Gilles Charvin

## Abstract

Automating the extraction of meaningful temporal information from sequences of microscopy images represents a major challenge to characterize dynamical biological processes. So far, strong limitations in the ability to quantitatively analyze single-cell trajectories have prevented large-scale investigations to assess the dynamics of entry into replicative senescence in yeast. Here, we have developed DetecDiv, a microfluidic-based image acquisition platform combined with deep learning-based software for high-throughput single-cell division tracking. We show that DetecDiv can automatically reconstruct cellular replicative lifespans with high accuracy and performs similarly with various imaging platforms and geometries of microfluidic traps. In addition, this methodology provides comprehensive temporal cellular metrics using time-series classification and image semantic segmentation. Last, we show that this method can be further applied to automatically quantify the dynamics of cellular adaptation and the real-time cell survival upon exposure to environmental stress. Hence, this methodology provides an all-in-one toolbox for high-throughput phenotyping for cell cycle, stress response, and replicative lifespan assays.

## Introduction

Epigenetic processes that span several division cycles are ubiquitous in biology and underlie essential biological functions, such as cellular memory phenomena(Caudron and Barral 2013; Bheda et al. 2020; Kundu, Horn, and Peterson 2007), differentiation, and aging (Denoth-Lippuner, Julou, and Barral 2014; Janssens and Veenhoff 2016). In budding yeast, mother cells undergo about 20 to 30 asymmetric divisions before entering senescence and dying (Mortimer and Johnston 1959). Over the last decades, this simple unicellular has become a reference model for understanding the fundamental mechanisms that control longevity (Denoth-Lippuner, Julou, and Barral 2014; Janssens and Veenhoff 2016). Several independent mechanistic models have been proposed to explain entry into replicative senescence, including asymmetric accumulation of extrachromosomal rDNA circles (ERCs) (Sinclair and Guarente 1997), protein aggregates (Aguilaniu et al. 2003), signaling processes associated with loss of vacuole acidity (Hughes and Gottschling 2012), or loss of chromatin silencing (Pal and Tyler 2016). Classical replicative lifespan assays (RLS) by microdissection, combined with genetic perturbations, have been decisive in identifying and characterizing genetic factors and pathways that influence longevity in budding yeast (McCormick et al. 2015). Similarly, enrichment techniques of aged mother cells in a batch provided further understanding of the physiology of cellular senescence in this model organism (Lindstrom and Gottschling 2009; Janssens et al. 2015).

However, how the appearance of markers of aging is coordinated temporally and causally remains poorly understood (Dillin, Gottschling, and Nyström 2014; C. He, Zhou, and Kennedy 2018). In part, this is due to the difficulty of directly characterizing the sequence of events that constitute the senescence entry scenario: RLS assays by microdissection generally give no information other than the replicative age upon cell death; old cells enrichment techniques ignore the well-known large cell-cell variability in the progression to senescence, which may blur the sequence of individual cellular events.

Based on pioneering work in yeast (Ryley and Pereira-Smith 2006) and bacteria (Wang et al. 2010), the development of microfluidics-based mother cell traps has partially alleviated these limitations by allowing continuous observation of individual cell divisions and relevant fluorescent cellular markers under the microscope from birth to death (Lee et al. 2012; Xie et al. 2012; Fehrmann et al. 2013). In these studies, monitoring individual cells over time in a microfluidic device has demonstrated the unique potential to quantitatively characterize the heterogeneity in cellular dynamics during aging. Recent years have seen a wide diversification of microfluidic devices aimed at improving both experimental throughput and cell retention rates (Jo et al. 2015; Liu, Young, and Acar 2015), (Li et al. 2017)). These new developments have helped to highlight the existence of independent trajectories leading to cell death (Li et al. 2017; Morlot et al. 2019; Li et al. 2020) and to better understand the physiopathology of the senescent state (Neurohr et al. 2018).

However, the hype surrounding these emerging microfluidic techniques has so far masked a key problem associated with high-throughput time-lapse imaging, namely the difficulty of extracting quantitative information in an efficient and standardized manner due to the manual aspect of the analysis (Huberts et al. 2014). In theory, expanding the number of individual cell traps and chambers on a microfluidic system makes it possible to test the effect of a large number of genetic and/or environmental perturbations on aging. Yet, in practice, this is out of reach since lifespan analyses require manual division counting and frequent corrections in cell segmentation. This problem has largely limited the interest of the “arms race” observed in recent years for the temporal tracking of individual cells during aging. This has also made it very difficult to cross-validate the results obtained by different laboratories, which is yet essential to advance our understanding of the mechanisms involved in aging.

Fortunately, the rapid development of powerful deep learning-based image processing methods in biology using convolutional neural networks (CNN) (Laine et al. 2021) suggests a way to overcome this important technical barrier. Recently, a study showed the potential of image classification by a CNN or a capsule network to classify the state of dividing yeast cells (*i*.*e*., budded, unbudded, etc.) trapped in a microfluidic device (Ghafari et al. 2021). However, due to the limited accuracy of the model, it has not demonstrated its ability to perform an automated division counting, let alone determine the RLS of individual cells. This is likely due to the fact that the misclassification of a single frame during the lifespan can dramatically compromise the accuracy of the RLS measurement.

Here, we report the development of DetecDiv, an integrated platform that combines high-throughput observation of cell divisions using a microfluidic device, a simple benchtop image acquisition system, and a deep learning-based image processing software with several image classification frameworks. Using this methodology, one can accurately track successive cell divisions in an automated manner and reconstruct RLS without human intervention, saving between 2 and 3 orders of magnitude on the analysis time. By combining this pipeline with additional deep-learning models for time-series classification and semantic segmentation, we provide a comprehensive toolset for an in-depth quantification of single-cell trajectories (*i*.*e*. division rate, mortality, size, and fluorescence) during entry into senescence and adaptation to environmental stress.

## Results

### An image sequence classification model for automated division counting and lifespan reconstruction

The primary scope of our present study was to overcome the current limitations inherent to the analysis of large-scale replicative lifespan assays by taking advantage of deep-learning image processing methods. Yet, we took this opportunity to provide improvements to individual mother cell trapping devices, in order to maximize the robustness of RLS data acquisition. Based on a design similar to that reported in previous studies (Jo et al. 2015; Liu, Young, and Acar 2015; Crane et al. 2014), we added small jaws on the top of the trap to better retain the mother cells in the traps (especially the old ones) (Figure 1 and Figure 1 - figure supplement 1G). In addition, we reduced the wall thickness of the traps to facilitate their deformation and thus avoid strong mechanical constraints when the cells become too big (Figure 1 - figure supplement 1D,G and supplementary text for details). Finally, we added a microfluidic barrier that filters cells coming from microcolonies located upstream of the trap matrix, which eventually clog the device and thus compromise the experiment after typically 24h of culture. Altogether, the microfluidic device features 16 independent chambers with 2000 traps each, eliciting multiple conditions and strains to be analyzed in parallel.

**Figure 1.**
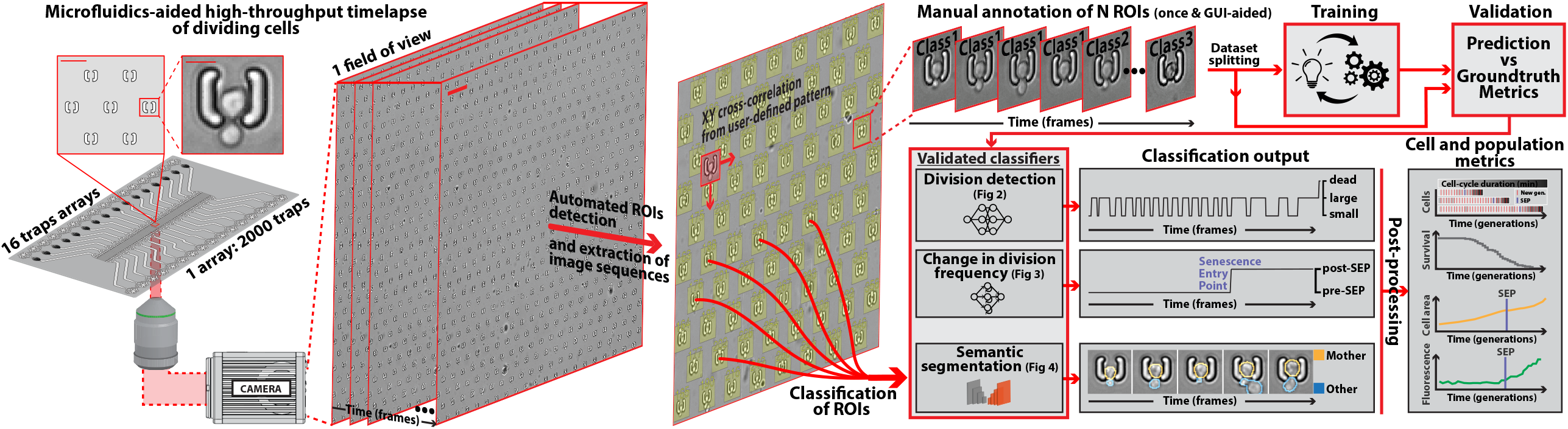
DetecDiv workflow. Left: Sketch of the hardware setup used to track divisions at the single-cell level. A microfluidic device, featuring 16 independent channels with 2000 individual cell traps in each (depicted with a zoom on the trap array (scale bar: 20μm) and zoom on one trap containing a budding yeast (scale bar: 5μm)), is imaged using timelapse microscopy. Middle-left: Typical temporal sequence of brightfield field of views obtained with the setup (scale bar: 60μm). Regions Of Interest (ROI) representing the traps are automatically detected using XY cross-correlation processing, and the temporal sequence of each ROI (trap) is extracted and saved. Top-right: Sketch of the training and validation pipeline of DetecDiv classifiers. A set of ROIs is picked from one (or several) experiments and annotated to form a ground-truth dataset. It is then split into a training set, used to train the corresponding classifier, and a test set used to validate the trained classifier. Bottom-right: Example of signals extracted from ROIs using DetecDiv classifiers. An image classifier can be used to extract oscillations of classes describing the size of the bud, from dividing cells, and thus the occurrence of new cell cycles (more details in Figure 2). A sequence classifier can be used to detect changes in cell-cycle frequency, such as a cell-cycle slowdown (Senescence Entry Point, SEP) (more details in Figure 4). A pixel classifier can be used to segment the mother cell from other cells, and from the background (more details in Figure 5). Using these classifiers on the same ROIs allows extracting quantitative metrics from dividing cells, at the single-cell and population level.

Next, we built a custom benchtop microscope (referred to as the “RAMM system” in the following, see methods for details) using simple optical parts to demonstrate that high-throughput division counting and quantitative RLS assays do not require any expensive fully-automated or high-magnification commercial microscopy systems. For this, we used a simple rigid frame with inverted epifluorescence optics, a fixed dual-band GFP/mCherry filter set, a bright field illumination column, a camera, and a motorized stage, for a total cost of fewer than 40k euros (Figure 1 - figure supplement 1A-B). Image acquisition, illumination, and stage control were all interfaced using the open-source Micromanager software (Edelstein et al. 2014). Using a 20x magnification objective, this “minimal” microscope allowed us to follow the successive divisions and the entry into senescence of typically 30000 individual cells in parallel with a 5-min resolution (knowing that there are ∼500 traps per field of view using the 20x objective).

This image acquisition system generates a large amount of cell division data (on the Terabytes scale depending on the number of channels, frames, and fields of view), only a tiny part of which can be manually curated in a reasonable time. In particular, the determination of replicative lifespans requires counting successive cell divisions until death, hence, reviewing all images acquired for each cell in each field of view over time. In addition, automating the division counting process is complicated by the heterogeneity in cell fate (*i*.*e*. cell-cycle durations and cell shape), especially during the entry into senescence.

To overcome this limitation, we have developed an image classification pipeline to count successive generations and reconstruct the entire lifespan of individual cells dividing in the traps (Figure 2A). For this, we have trained a convolutional neural network (CNN) based on the “Inception” architecture (Szegedy et al. 2015) to predict the budding state of the trapped cells by assigning one of six possible classes (unbudded, small-budded, large-budded, dead, empty trap, and clogged trap) to each frame (Figure 2A, Top). In this framework, the alternation between the ‘large budded’ or ‘unbudded’ and the ‘small budded’ states reveals bud emergences. The cell cycle durations can be deduced by measuring the time interval between successive budding events, and the occurrence of the ‘dead’ class determines the end of the cell’s lifespan (Figure 2A, Bottom). We selected this classification scheme - namely, the prediction of the budding state of the cell - over the direct assessment of cell division or budding (e.g., “division” versus “no division”) because division and budding events can only be assessed by comparing successive frames, which is impossible using a classical CNN architecture dedicated to image classification, which takes a single frame as input.

**Figure 2.**
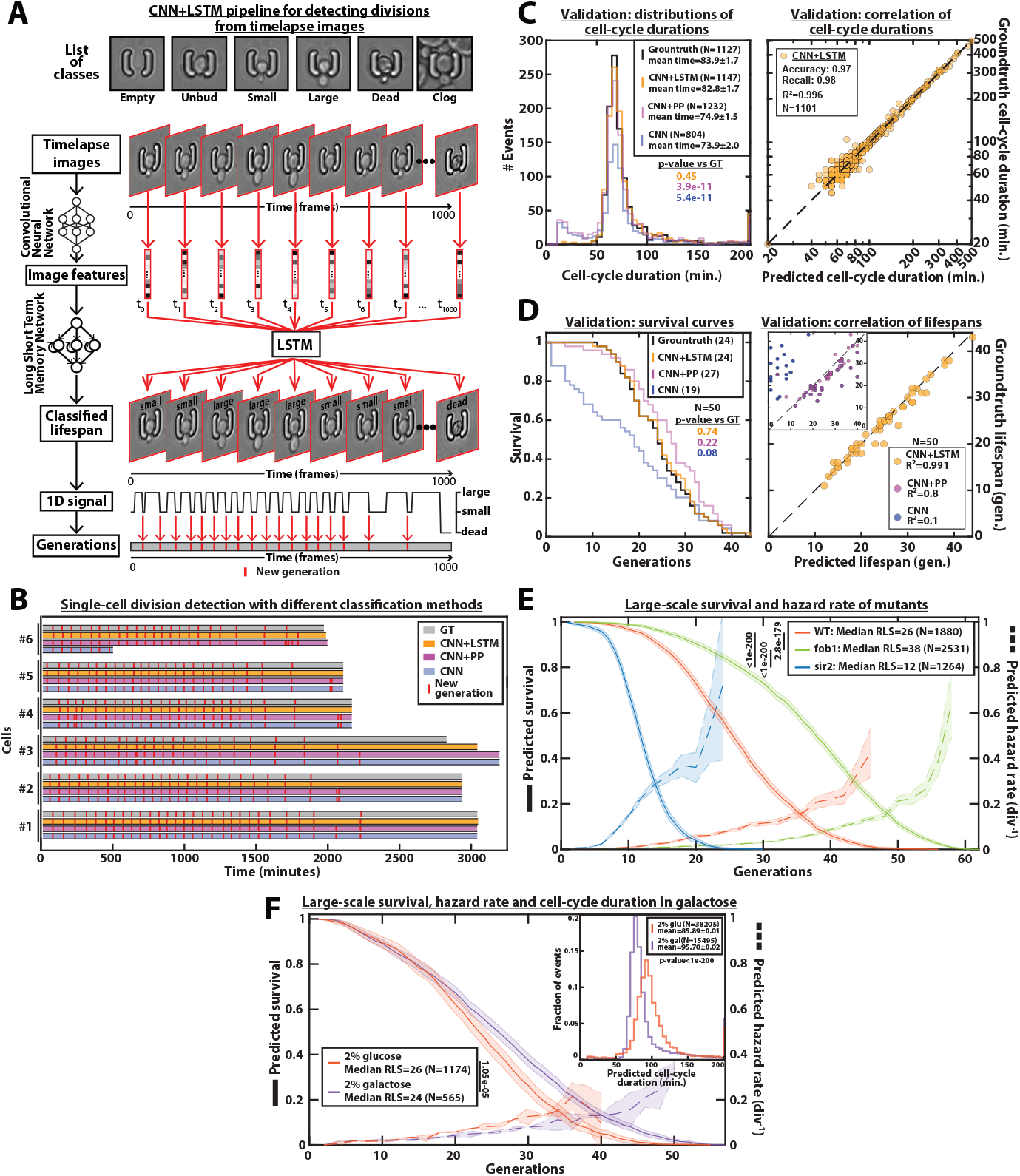
DetecDiv cell-cycle duration predictions and RLS reconstruction pipeline. A. Principles of the DetecDiv division tracking and lifespan reconstruction pipeline; Brightfield images are processed by a convolutional neural network (CNN) to extract representative image features. The sequence of image features is then processed by a long short-term memory network (LSTM) that assigns one of the 6 predefined classes (‘unbud’, ‘small’, ‘large’, ‘dead’, ‘clog’, ‘empty’), taking into account the time dependencies. Temporal oscillations between “large” and “small” or “large” and “unbudded” indicate the beginning of a new generation (*i*.*e*. cell-cycle). The appearance of the “dead” class marks the end of the lifespan. B. Comparison of the different methods used for 6 sample cells. The gray bars represent the ground truth data made from manually annotated image sequences. Colored lines indicate the corresponding predictions made by CNN/LSTM (orange), the CNN + post-processing (magenta), and the CNN (blue) networks (see Methods and supplementary text for details). The red segments indicate the position of new generation events. C. Left: histogram of cell-cycle durations representing ground truth data and predictions using different processing pipelines. The p-value indicates the results of a rank-sum test comparing the predictions to the ground truth for the different pipeline variants. The total number of generations annotated in the ground truth or detected by the networks is indicated in the legend. Right: Scatter plot in log scale representing the correlation between ground truth-calculated cell-cycle durations and those predicted by the CNN/LSTM network. R^2^ represents the coefficient of correlation between the two datasets. Accuracy and recall are defined in the supplementary text. D. Left: cumulative distribution showing the survival of cells as a function of the number of divisions (N=50 cells). The numbers in the legend indicate the median replicative lifespans. The p-value indicates the results from a statistical rank-sum test. Right: Scatter plot representing the correlation of the replicative lifespans of individual cells obtained from the ground truth with that predicted by the CNN/LSTM architecture (N=50). Inset: same as the main plot, but for the CNN and CNN with post-processing pipelines. R^2^ indicates the coefficient of correlation between the two datasets. E. Replicative lifespans obtained using the CNN/LSTM network for longevity mutants (solid colored lines, genotype indicated). The shading represents the 95% confidence interval calculated using the Greenwood method (Pokhrel, Dyba, and Hakulinen 2008). The median RLS and the number of cells analyzed are indicated in the legend. The dashed lines with shading represent the hazard rate and its standard deviation estimated with a bootstrap test (N=100). Results from log-rank tests (comparing WT and mutant distributions) are indicated on the left of the legend. F. Same as E) but for WT cells grown in 2% glucose or 2% galactose (colored lines). Inset: Same as C) (Left) but with the conditions from the main.

To train and evaluate the performance of the classifier, we generated a manually annotated dataset (referred to as “ground truth” in the following) by arbitrarily selecting 250 traps containing situations representative of all cellular states from different fields of view and independent experimental replicates. To do so, we created a graphical user interface (GUI) to automatically detect the traps (by image auto-correlation with a user-selected reference pattern) and extract a 4-dimensional matrix (x,y, channel, time) for each trap (referred to as a Region Of Interest, or ROI, in the following, see Figure 1). Then, the GUI was designed to assist the users with rapid screening and assignment of successive individual frames using keyboard shortcuts. Using this tool, it took about 3-9min to annotate a series of 1000 frames for a single ROI. Hence, the entire ground truth dataset was generated within 15-20h of work. We then arbitrarily split the ground truth into two independent datasets: a training dataset with 200 ROIs was used to train the model using classical stochastic gradient descent (see Methods for details) and a separate test dataset containing 50 ROIs was used to benchmark the performance of the classifier after training (as summarized in Figure 1).

Benchmarking the classifier consisted of three steps: first, we computed the confusion matrices (Figure 2 - figure supplement 2A) as well as the classical metrics of accuracy, recall, and F_1_-score. The F_1_-score was found to be higher than 85% for all classes (Figure 2 - figure supplement 2C). Next, the predictions of budding events were compared to the manually annotated data. Despite a good visual match between the ground-truth and the CNN predictions, the distribution of division times revealed that the model tends to predict “ghost” divisions of abnormally short duration (Figure 2B). In addition, sporadic misclassification could falsely assign a cell to the “dead” state, thus decreasing the number of total generations predicted based on the test dataset (N=1127 for the groundtruth versus N=804 for the CNN model, see Figure 2C). Last, by comparing the lifespan predictions to the corresponding groundtruth data, we observed a striking underestimate of the overall survival (Figure 2D), due to sporadic misassignments in the “dead” class (Figure 2 - figure supplement 1B).

These problems could be partially alleviated by post-processing the predictions made by the CNN (see “CNN+PP” in Figure 2B-D and supplementary text for details). Indeed, by ignoring isolated frames with a “dead” class, we could greatly reduce the number of cases with premature cell death prediction, yet we failed to efficiently remove ghost divisions, hence leading to an overestimate of the RLS and a large number of short cell-cycles (Figure 2C-D).

An inherent limitation to this approach is that images are individually processed without taking the temporal context into account. Although a more complex post-processing routine could be designed to improve the robustness of the predictions, it would come at the expense of adding more *ad hoc* parameters, hence decreasing the generality of the method. Therefore, to circumvent this problem, we decided to combine the CNN image classification with a long short-term memory network (LSTM) (Venugopalan et al. 2015; Hochreiter and Schmidhuber 1997), to take into account the time-dependencies between images (Figure 2A, Middle). In this framework, the CNN was first trained on the individual images taken from the training set similarly as above. Then, the CNN network activations computed from the temporal sequences of images were used as input to train an LSTM network (see suppl. for details). Following this training procedure, the assembled CNN + LSTM network was then benchmarked similarly as described above. We obtained only a marginal increase in the classification metrics compared to the CNN network (about 90-95% precision and recall for all classes, see Figure 2 - figure supplement 2A-B). Yet, strikingly, the quantification of division times and cellular lifespan both revealed considerable improvements in the accuracy: “ghost” divisions were drastically reduced if not completely removed, the distribution of cell-cycle duration was indistinguishable from that of the ground truth (p=0.45, Figure 2C), and the difference between the number of divisions predicted by the network and the actual number was less than 2% (N=1147 and N=1127, respectively, see left panel on Figure 2C). In addition, the Pearson correlation coefficient for ground truth vs prediction was very high (R^2^=0.996, see right panel on Figure 2C). This indicates that mild classification errors may be buffered and hence do not affect the accuracy in the measurements of cell cycle durations. Moreover, it suggests that the network was robust enough to ignore the budding status of the daughter cells surrounding the mother cell of interest (Figure 2 - figure supplement 3). Similarly, the predicted survival curve was almost identical to that computed from the ground truth (p=0.74, Figure 2D and Movie M1) and the corresponding Pearson correlation reached 0.991 (vs 0.8 and 0.1 for the CNN+PP and CNN, respectively). Altogether, these benchmarks indicated that only the combined CNN+LSTM architecture provided the necessary robustness to provide an accurate prediction of individual cellular lifespan based on image sequence classification.

Following its validation, we deployed this model to classify all the ROIs from several fields of view extracted from 3 independent experiments. We were thus able to obtain a survival curve with N=1880 lifespans in a remarkably short time (Figure 2E): less than 3.5s were necessary to classify 1000 images using a Tesla K80 GPU (*i*.*e*. more than 100 times faster than manual annotation). Conversely, it would have taken 130 days working 24 hours a day for a human being to generate this plot by manual annotation. To further apply the classification model trained on images of wild-type (WT) cells, we measured the large-scale RLS in two classical longevity mutants. Remarkably, we recapitulated the increase (resp. decrease) in longevity observed in the *fob1Δ* (resp. *sir2Δ)* mutant (Defossez et al. 1999; Lin, Defossez, and Guarente 2000) and we could compute the related death rate with a high-confidence interval thanks to this large-scale dataset (Figure 2E). In addition, we performed comparative measurements of cell-cycle duration (N=38205 events and N= 15495 events for glucose and galactose, see Figure 2F inset) and RLS (median=26 generations (N=1174 events) and median=24 (N=565 events) for glucose and galactose, respectively, Figure 2F) using glucose and galactose as carbon sources. Our results were in line with previous measurements (Liu, Young, and Acar 2015; Frenk et al. 2017) and hence further confirmed the validity of our analysis pipeline. Therefore, despite a significant initial investment (*i*.*e*. ∼ one week of work) to generate the training and test datasets (250 ROIs in this case), our study shows that the classification pipeline subsequently saves months of work compared to manual analysis of the data.

### Application of the Detecdiv pipeline to different imaging platforms and microfluidic device geometries

To test the robustness and versatility of our analysis pipeline, we proceeded to the analysis of several datasets obtained under various conditions. First, we performed experiments with the same microfluidic system but using a commercial microscope with 60x magnification. Following the training of the classifier on 80 ROIs and testing on 40 independent ROIs, we observed similar results to those obtained with the RAMM system and a 20x objective (compare the “specialist” columns for the panels in Figure 3A and 3B): the classification benchmarks were greater than 90%, the error rate on the number of divisions detected was a few percents, and the cell-cycle length distributions were similar between prediction and ground truth. This first demonstrated that neither the RAMM imaging system nor the 20x magnification are required to guarantee successful division counting and lifespan reconstruction with our analysis pipeline.

**Figure 3.**
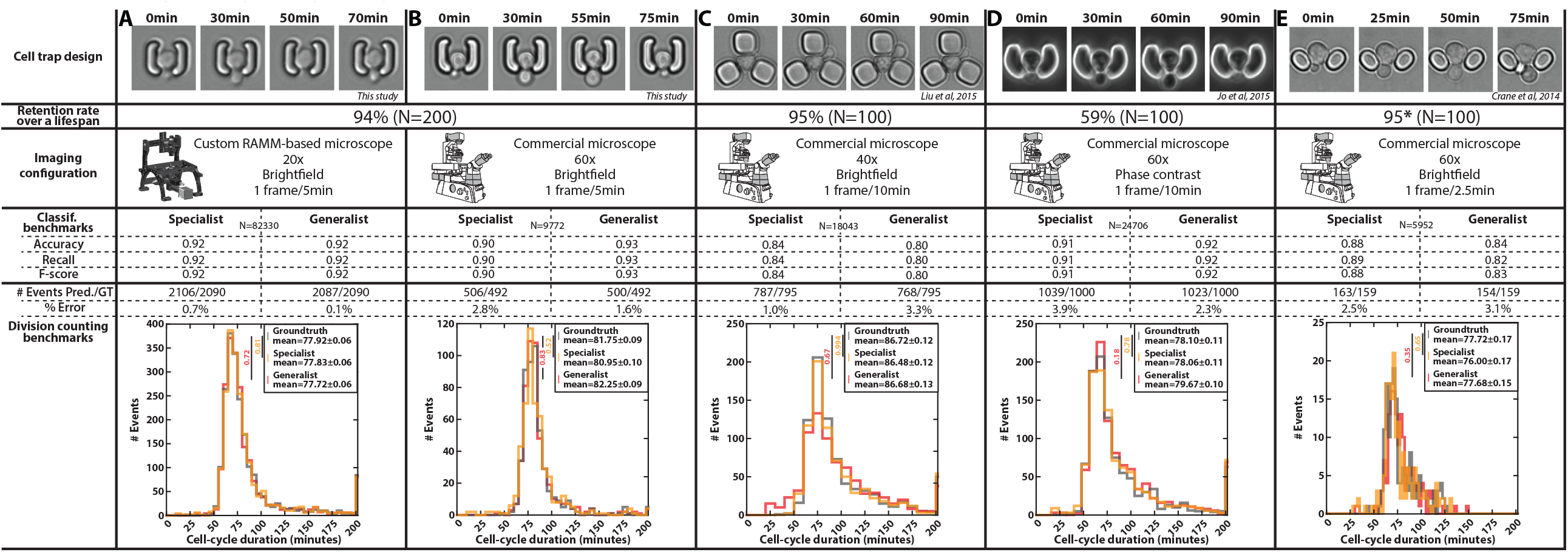
Automated detection of divisions with images from different microfluidic devices and imaging setups. Classification benchmarks and performances of the divison detection of a CNN+LSTM image classifier, trained on time-lapses of cells from different sources. A specialist classifier was trained independently for each source, while a generalist classifier was trained on a mixed dataset generated from all the sources. A. Cell trap and imaging setup developed in this study, with a framerate of 1 frame/5min. B. Cell trap developed in this study imaged with a 60x objective mounted on a commercial imaging system with a framerate of 1 frame/5min. C. Cell trap from the Acar lab (Liu, Young, and Acar 2015) imaged with a 40x objective mounted on a commercial imaging system with a framerate of 1 frame/10min. D. Cup-shaped trap similar to Jo et al. 2015 (Jo et al. 2015), imaged with a 60x phase-constrast objective mounted on a commercial imaging system with a framerate of 1 frame/10min. E. Cell trap from the Swain lab (Crane et al. 2014) imaged with a 60x objective mounted on a commercial imaging system with a framerate of 1 frame/2.5min.

In addition, we gathered time-lapse microscopy datasets from several laboratories using microfluidic cell trapping systems with different geometries and based on acquisition under various imaging conditions (Figures 3C and 3E) (Crane et al. 2014; Liu, Young, and Acar 2015). We also included data generated in our lab based on a device similar to that used in Jo et al.(Jo et al. 2015) (Figure 3D). For each trap geometry, we manually evaluated the retention rate of a mother cell during a lifespan. Indeed, high retention is key to getting a reliable measurement of the RLS (*i*.*e*., to ensure that mother cells are not eventually replaced by their daughters). This analysis revealed that a “semi-open” geometry (as in the design shown in Figure 3D, Figure 1 - figure supplement 1G, and Figure 3 - figure supplement 1) did not prevent large mother cells from being sporadically expelled during the budding of their daughters, unlike other cell trap shapes. Of note, the geometry proposed by Crane et al.(Crane et al. 2014) (Figure 3E) was not tested on an entire lifespan, but only on about 7 generations, hence leading to an overestimation of the retention rate (it was reported to be below 50% in the original paper).

For each dataset, we trained a specific classifier (or “specialist”) on 80 ROIs and validated it on 40 independent ROIs. The different benchmarks (*i*.*e*., classification performance and division predictions) showed that each specialist performed very well on each specific test dataset, thus confirming further that our analysis pipeline is robust and applicable to different cell trapping configurations.

Last, instead of training the classifiers separately on each dataset, we asked whether a unique classifier would have sufficient capacity to handle the pooled dataset with all imaging conditions and trap geometries used in Figure 3. Strikingly, this “generalist” model showed comparable performance to the different specialists. This approach thus further highlighted the versatility of our methodology and demonstrated the interest in aggregating data sets to ultimately build a standardized reference model for counting divisions, independently of the specific imaging conditions.

### Automated quantification of cellular physiological decline upon entry into senescence

Aging yeast cells have long been reported to undergo a cell division slowdown when approaching senescence (Mortimer and Johnston 1959), a phenomenon that we have since quantified and referred to as the <u>S</u>enescence <u>E</u>ntry <u>P</u>oint or SEP (Fehrmann et al. 2013). More recently, we have demonstrated that this quite abrupt physiological decline in the cellular lifespan is concomitant with the accumulation of extrachromosomal rDNA circles (ERCs) (Morlot et al. 2019), a long described marker of aging in yeast (Sinclair and Guarente 1997). Therefore, precise identification of the turning point from healthy to pathological state (named pre-SEP and post-SEP in the following, respectively) is essential to capture the dynamics of entry into senescence, and even more so since the large cell-cell variability in cell death makes trajectory alignment from cell birth irrelevant(Fehrmann et al. 2013; Morlot et al. 2019). To achieve this analysis in an automated manner, we sought to develop an additional classification scheme as follows: we trained a simple LSTM sequence-to-sequence classifier to assign a ‘pre-SEP’ or ‘post-SEP’ label (before or after the SEP, respectively) to each frame, using the sequence of cellular state probabilities (*i*.*e*., the output of the CNN+LSTM image classifier described in Figure 2A) as input (Figure 4A). The ground truth was generated by visual inspection using a graphical user interface representing the budding status of a given cell over time. Same as above, we used 200 manually annotated ROIs for the training procedure and reserved 47 additional ones that were never “seen” by the network to evaluate the predictions. Comparing the predictions to the groundtruth revealed that we could successfully identify the transition to a slow division mode (R^2^=0.93, see Figure 4B-C and Figure 4 - figure supplement 1). Hence, we could recapitulate the rapid increase in the average cell-cycle durations after aligning individual trajectories from that transition (Figure 4D), as described before (Fehrmann et al. 2013). These results show that complementary classifiers can be used to process time series output by other classification models, allowing further exploitation of relevant dynamic information, such as the entry into senescence.

**Figure 4.**
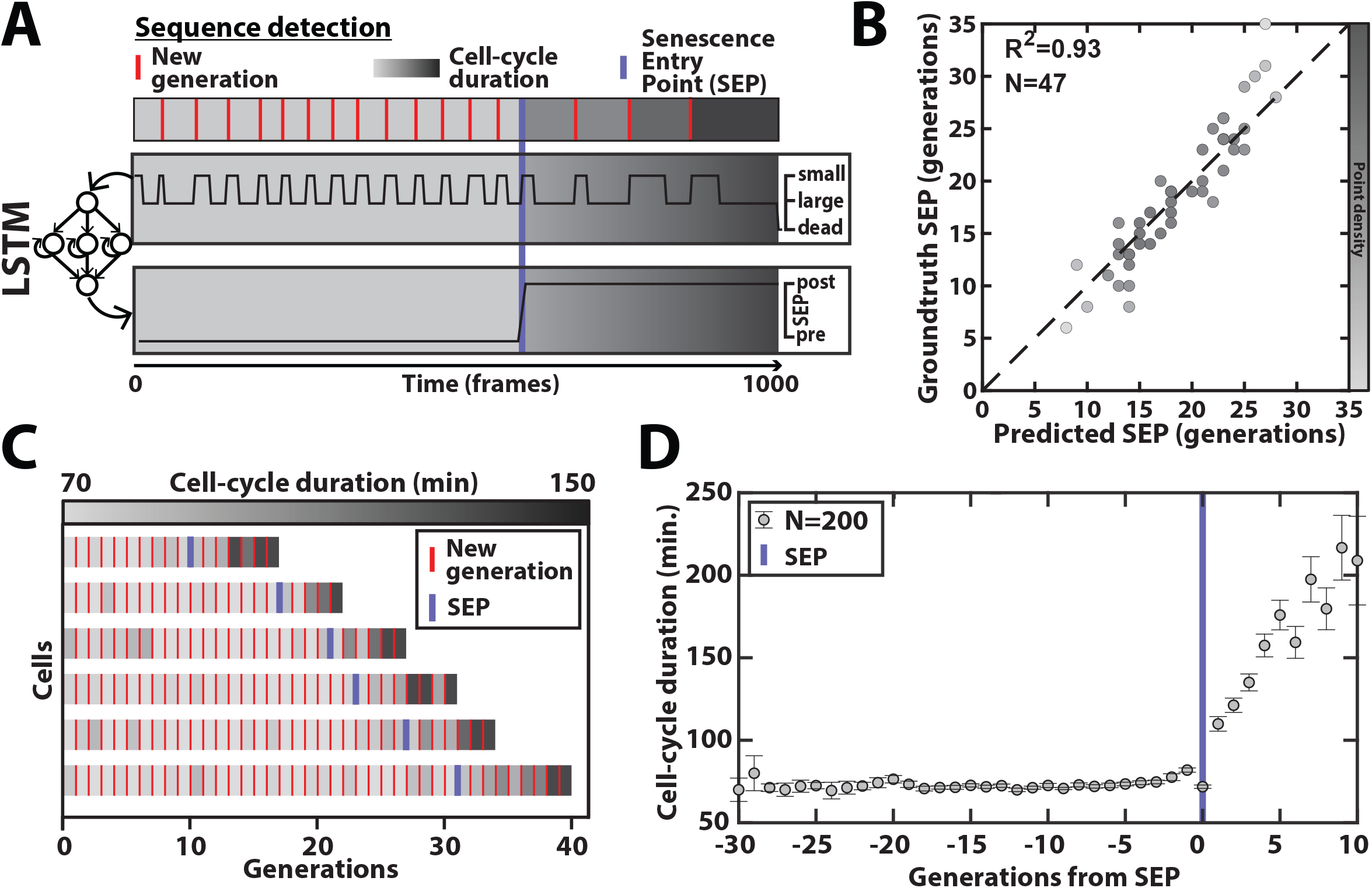
Deep learning-based measurement of the dynamics of entry into senescence. A. Sketch depicting the detection of the Senescence Entry Point (SEP). The temporal sequence of classes probabilities (ie, unbud, small, large, dead) is fed into an LSTM network that predicts the SEP by assigning one of the two predefined classes pre-SEP or post-SEP to each frame. B. Correlogram showing the correlation between the SEP predicted by the LSTM network and the ground truth data, obtained as previously described (Fehrmann et al. 2013). The color code indicates the local density of the points using arbitrary units as indicated by the color bar. C. Sample trajectories indicating the successive divisions of individual cells (red lines) along with the division times (color-coded as indicated). D. Average cell-cycle duration versus generation index after aligning all individual trajectories from the SEP (Fehrmann et al. 2013). Each point represents an average over up to 200 cell trajectories. The error bar represents the standard error-on-mean.

### Cell contour determination and fluorescence image quantification by semantic segmentation

Quantifying the dynamics of successive divisions is an indispensable prerequisite for capturing phenomena that span multiple divisions such as replicative aging. However, in order to make the most of the possibilities offered by photonic microscopy, it is necessary to develop complementary cytometry tools. For this purpose, semantic segmentation (based on the classification of pixels according to a finite number of classes) has seen a growing interest recently to process biomedical images, from the pioneering development of the U-Net architecture (Ronneberger, Fischer, and Brox 2015) to more sophisticated versions allowing the segmentation of objects with low contrast and/or in dense environments (Stringer et al. 2021). In addition, specific implementations of U-Net have demonstrated the broad potential of this architecture for segmenting (Dietler et al. 2020) and tracking (Lugagne, Lin, and Dunlop 2020; Ershov et al. 2021) cells in various model organisms.

Here, we have implemented an encoder/decoder network based on a Resnet50 CNN (K. He et al. 2016) and the DeepLab v3+ architecture (Chen et al. 2018), (Figure 5 - figure supplement 1), to segment brightfield images (Figure 5A, Movie M2, and supplementary text). We have trained the model on ∼1400 manually segmented brightfield images using three output classes (*i*.*e*., ‘background,’ ‘mother cell,’ ‘other cell’) in order to discriminate the mother cell of interest from the surrounding cellular objects. We have used a separate test dataset containing ∼500 labeled images to evaluate the performance of the classifier. To generate the ground truth data required to feed both the training and test datasets, we have developed a graphical user-interfaced routine to “paint” the input images, a process which took about 15-30 seconds per image depending on the number of cells in a 60×60 pixel-large field of view. Our results revealed that mother cells contours could be determined accurately with a trained classifier (Figure 5A-C and Figure 5 - figure supplement 2A-D). In addition, we used a cross-validation procedure based on random partitioning of training and test datasets that highlighted the robustness of the classification (Figure 5 - figure supplement 2E). Overall, this segmentation procedure allowed us to quantify the dynamics of volume increase of the mother cell (Figure 5C-D), as previously reported (Morlot et al. 2019).

**Figure 5.**
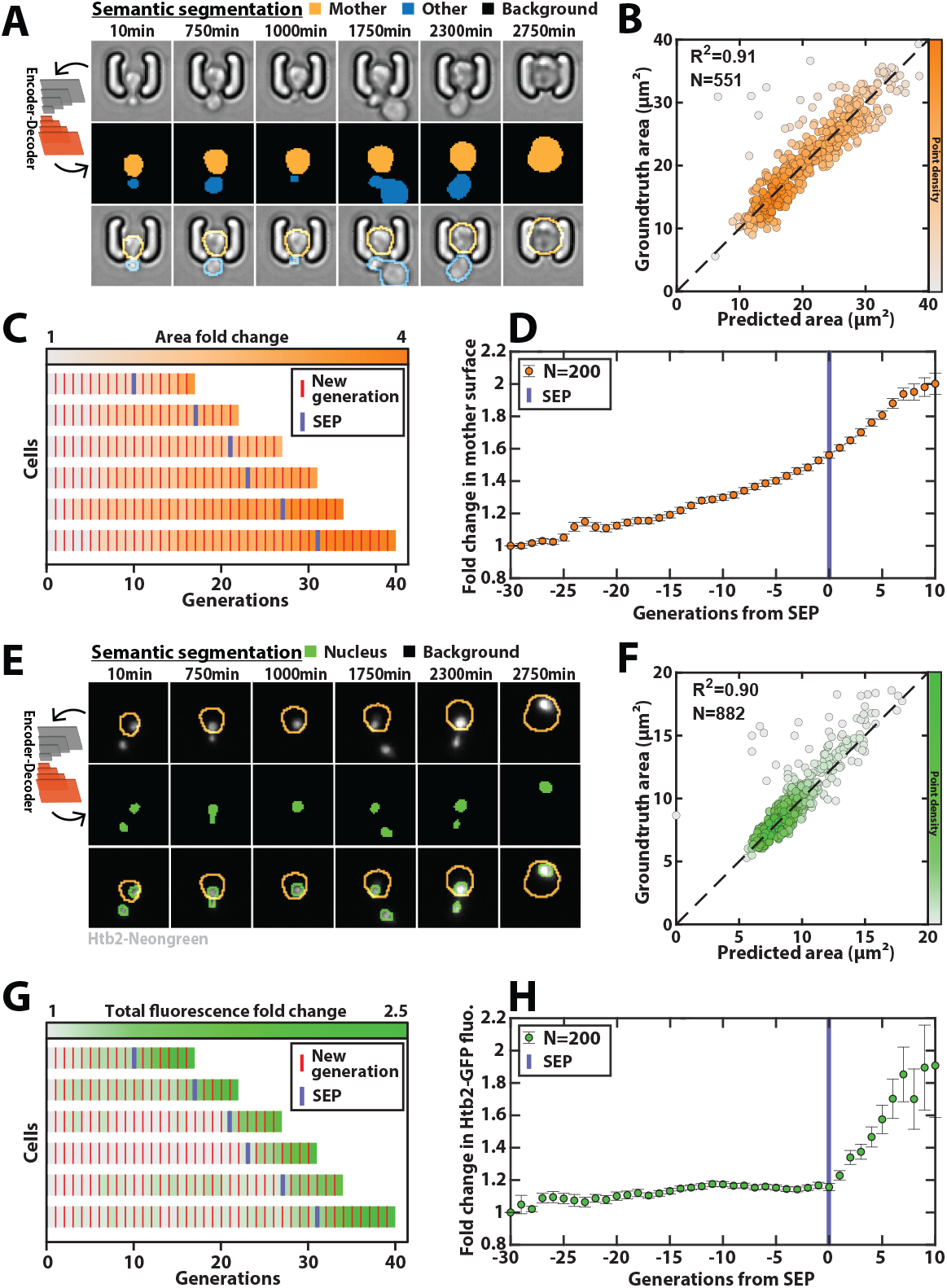
Deep learning-based semantic segmentation of cells and nuclei. A. Principles of semantic cell contours segmentation based on brightfield images; Top and middle row) Individual brightfield images were processed by the DeeplabV3+ network that was trained to perform pixel classification using three predefined classes representing the background (black), the mother cell of interest (orange), or any other cell in the image (blue). Bottom row: overlay of brightfield images with segmented cellular contours. B. Correlogram showing the correlation between individual cell area predicted by the segmentation pipeline and the ground truth data, obtained by manual annotation of the images. The color code indicates the local density of the points using arbitrary units. C. Sample trajectories indicating the successive generations of individual cells (red lines) along with the cell surface area (color-coded as indicated). D. Average mother cell surface area versus generation index after aligning all individual trajectories from the SEP (Fehrmann et al. 2013). Each point represents an average over up to 200 cell trajectories. The error bar represents the standard error-on-mean. E. Principles of semantic cell nuclei segmentation based on fluorescent images of cells expressing a histone-Neongreen fusion. The semantic segmentation network was trained to classify pixels between two predefined classes (‘background’ in black, ‘nucleus’ in green). F. Same as B) but for nuclear surface area. G. Same as C) but for total nuclear fluorescence H. Same as in D) but for total nuclear fluorescence

Last, a similar training procedure with ∼3000 fluorescence images with a nuclear marker (using a strain carrying a histone-Neongreen fusion) yielded accurate nuclei contours (Figure 5E-F, Figure 5 - figure supplement 3). It successfully recapitulated the sharp burst in nuclear fluorescence that follows the Senescence Entry Point (Figure 5G-H) (Morlot et al. 2019).

### Automated quantitative measurements of the physiological adaptation to hydrogen peroxide

Beyond replicative longevity analyses, we wondered if this automated pipeline could be applied to other biological contexts, in which cell proliferation and cell death need to be accurately quantified over time. Hence, we sought to measure the dynamics of the physiological adaptation of yeast cells subjected to hydrogen peroxide stress.

For this purpose, young cells were abruptly exposed to different stress concentrations, ranging from 0 to 0.8mM H_2_O_2_, and observed over about 15h (Figure 6A). We used a strain carrying the Tsa1-GFP fusion protein (*TSA1* encodes a peroxiredoxin, a major cytosolic antioxidant overexpressed in response to oxidative stress) as a fluorescent reporter of the cellular response to this stress (Goulev et al. 2017).

**Figure 6.**
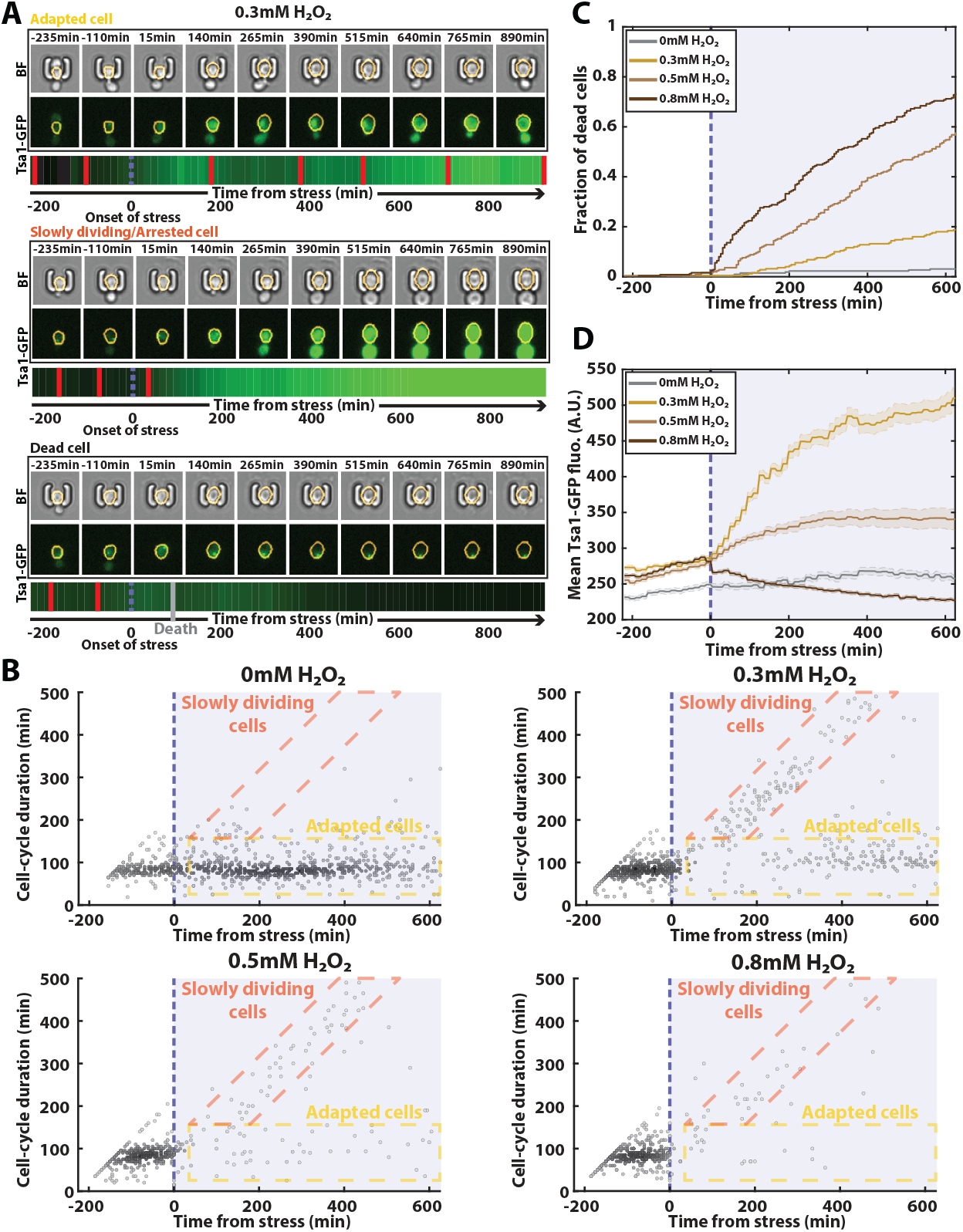
Automated analysis of the stress response to H_2_O_2_ using DetecDiv. A. Successive brightfield and Tsa1-GFP images of three representative cells submitted to 0.3mM of H_2_O_2_ and corresponding to a different fate. The orange contour of the cell is determined using the segmentation described in figure 5, and the total GFP fluorescence inside it is depicted as a function of time, where red bars indicates a new generation and the purple dotted bar indicated the onset of H_2_O_2_. B. Scatter plot of automatically detected cell-cycle durations versus time of 500 cells submitted to different doses of H_2_O_2_. The purple area indicates the presence of the indicated dose of H_2_O_2_. Darker dots indicate higher point density. C. Fraction of dead cells versus time as automatically detected by the CNN+LSTM classifier, under different H_2_O_2_ doses. The purple area indicates the presence of the indicated dose of H_2_O_2_. D. Mean Tsa1-GFP fluorescence from cells submitted to different doses of H_2_O_2_. The purple area indicates the presence of the indicated dose of H_2_O_2_.

In this context, we first sought to characterize the dynamics of the cell-cycle by using the classifier reported in Figure 2 - without doing any retraining - to detect divisions during the experiment (using N=250 ROIs). Our automated analysis revealed different possible cell fates, whose proportions varied according to the stress concentration (Figure 6A and 6B): in the absence of stress (0mM), cells maintained a constant division rate throughout the experiment; in contrast, at 0.3mM, the population partitioned between cells that recovered a normal division rate after experiencing a delay (see the “adapted cells” in Figure 6B) and others that seemed unable to resume a normal cell-cycle (see the “slowly dividing cells” in Figure 6B), in agreement with previous results (Goulev et al. 2017).

Higher doses of stress (0.5mM and 0.8mM) saw these populations gradually disappear, indicating that very few divisions occur at these elevated doses. To check this further, we exploited further the outputs of the classifier to score the onset of cell death for each trapped cell (Figures 6A and 6C). Our analysis revealed a progressive, dose-dependent increase in the fraction of dead cells over time. These results thus demonstrated the possibility to perform real-time and quantitative measurement of the cell death rate in response to an environmental insult, which is rarely precisely done due to the difficulty of precisely scoring dead cells in a population of cells without any additional viability marker.

Finally, we used our semantic segmentation model (reported in Figure 5) to quantify cytoplasmic fluorescence over time in the stress response experiment (Figure 6A). The population-averaged mean cytoplasmic fluorescence revealed a significant increase at 0.3mM H_2_O_2_ due to the transcriptional upregulation of antioxidant genes, as previously described (Goulev et al. 2017). However, this average upregulation of Tsa1 was lessened at higher doses, an effect we attribute to the large fraction of cell death observed in these conditions (e.g.: bottom cell in Figure 6A). Altogether, these results indicate that DetecDiv used with single-cell traps provides a highly suitable method for quantifying both cell division rate and mortality in real-time in a controlled environment.

## Discussion

In this study, we have developed a pipeline based on the combined use of two architectures, namely a CNN+LSTM network for the exploitation of temporal information and semantic segmentation (DeepLabV3+) for the quantification of spatial information. We demonstrate that it can successfully characterize the dynamics of multi-generational phenomena, using the example of the entry into replicative senescence in yeast, a difficult case study that has long resisted any automated analysis. We also successfully used our classification model to score cellular adaptation and mortality in the context of the physiological stress response to hydrogen peroxide. We envision that this methodology will unleash the potential of microfluidic cell trapping devices to quantify temporal physiological metrics in a high-throughput and single-cell manner.

The major novelty of this work lies in the development of an analysis method to automatically obtain survival curves and cytometric measurements during the entry into senescence from raw image sequences. Nevertheless, we also focused our efforts on improving traps to increase the efficiency of RLS assays in microfluidics. Also, we have built a minimal optical system (yet with a motorized stage) assembled from simple optical components (*i*.*e*., no filter wheel, fixed objective), for a price of about one-third that of a commercial automated microscope, which can be made accessible to a larger community of researchers. Although many laboratories use common imaging platforms with shared equipment, it is important to note that the cost of an hour of microscopy is around 10-20 euros in an imaging facility. As an RLS assay typically lasts 60-80 hours, these experiments may not be affordable. Developing a simple system can therefore quickly pay off if the lab does not have its own microscope.

Using this experimental setup, we showed that our analysis pipeline works perfectly even with a low optical resolution (i.e. the theoretical resolution of our imaging system with a 20x, N.A. 0.45 objective is ∼0.7microns), and without any contrast-enhancing method. In practice, it might be desirable for some applications to use higher magnification to better preserve spatial information and analyze the detailed localization of fluorescent markers. Yet, using the same microfluidic device described here, we showed that DetecDiv works similarly with higher magnification objectives and different imaging systems. In addition, we demonstrated that division detection can also be performed with cells growing in traps with different geometries (Figure 3, Figure 1 - figure supplement 1G, and Figure 3 - figure supplement 1). Furthermore, a unique classifier trained on a large collection of images obtained under broad imaging contexts can also achieve accurate division detection. This may be instrumental to standardize the quantitative analysis of replicative lifespan data in the yeast aging community.

However, one limitation to applying our analysis pipeline with a broad range of trap geometries is that the accuracy of RLS measurements may be affected when using designs with a low retention rate. Although lifespan trajectories can be marked as ‘censored’ when the mother cell leaves the traps (as proposed in a recently released preprint (<u>Thayer et al. 2022</u>)), our method is currently unable to systematically detect when a mother cell is replaced by its daughter (e.g. cell traps in Figure 3D). Therefore, we believe that retention is an essential feature to consider when designing the geometry of a trap.

An important advantage of individual cell trapping is that it makes image analysis much simpler than using chambers filled with two-dimensional cell microcolonies. Indeed, individual traps behave as a “hardware-based cell tracking” method, thus alleviating the need to identify and track objects spatially, a procedure that provides an additional source of errors. Because the cells of interest are located in the middle of the traps, the learning process can focus the attention of the classifier on the state of the mother cell only (e.g. small-budded, large-budded, etc.), hence the specific state of the few cells surrounding it may not influence the reliability of the classification of the mother (Figure 2 - figure supplement 3 for specific examples). In addition, a powerful feature of whole image classification is that it can easily be coupled to a recurrent neural network (such as an LSTM network), thus opening the door to more accurate analyses that exploit temporal dependencies between images, as demonstrated in our study.

Beyond the tracking of successive divisions, complementary methods are necessary to characterize the evolution of cell physiology over time. In our study, we used semantic segmentation to delineate the contours of cell bodies over time. Same as above, the ability to discriminate the mother cell of interest from the surrounding cells results is facilitated by the conserved position of the mother at the center of the trap. However, a limitation of our classification scheme is that the buds that arise from the mother cell can not be identified, and further work is necessary to assess the requirements (e.g. the size of the training set) to achieve a successful segmentation using a separate ‘bud’ class. Thus, it is currently impossible to measure the growth rate (in volume) of the mother cell over time (most of the biomass created during the budded period goes into the bud) and it precludes analyzing fluorescent markers that would localize into the bud. Future work may explore how the use of additional segmentation classes or the use of tracking methods could complement our pipeline to alleviate this limitation. Alternatively, the development of an instance segmentation method (<u>He et al. 2017; Prangemeier et al. 2022</u>) could also facilitate the identification and separation of different cell bodies in the image.

Unlike classical image analysis methods, which require complex parameterization and are highly dependent on the problem being addressed, the well-known advantage of machine learning is the versatility of the models, which can be used for a variety of tasks. Here, we show that our division counting/lifespan reconstruction classifier can be used right away to quantify cellular dynamics and mortality in response to hydrogen peroxide stress. We envision that DetecDiv could be further applied in different contexts without additional development - yet with potential retraining of the classifier with complementary data, and/or following the definition of new classes. For example, it could be useful to develop a classifier able to identify different cell fates during aging based on image sequences (e.g. *petite* cells (Fehrmann et al. 2013), or mode 1 versus mode 2 aging trajectories (Jin et al. 2019)), as well as during induced (Bagamery et al. 2020) or undergone (Jacquel et al. 2021) metabolic changes. More generally, the rationalization of division rate measurements in a system where there is no competition between cells offers a unique framework to study the heterogeneity of cell behaviors in response to environmental changes (stress, chemical drugs, etc.), as demonstrated in our study and evidenced by the rise of high-throughput quantitative studies in bacteria (Bakshi et al. 2021). Mechanistic studies of the cell cycle could also benefit from a precise and standardized phenotypic characterization of the division dynamics. Along this line, beyond the classification models described in this study, we have integrated additional frameworks, such as image and image sequence regressions (Supplementary Table 2), which could be useful to score fluorescent markers quantitatively and over time (e.g. mitotic spindle length inference, scoring of the mitochondrial network shape, etc.). We envision that the kind of approach described here may be easily transferred to other cellular models to characterize heterogeneous and complex temporal patterns in biological signals.

## Methods

### Strains

All strains used in this study are congenic to S288C (Supplementary Table 1 for details). See supplementary methods for detailed protocols for cell culture.

### Microfabrication and microfluidics

The designs were created on AutoCAD to produce chrome photomasks (jd-photodata, UK). The microfluidic master molds were then made by standard photolithography processes (see supplementary text for details).

The microfluidic device is composed of geometric microstructures that allow mother cells trapping and flushing of successive daughter cells (Figure 1 - figure supplement 1 and supplementary text). The cell retention efficiency of the traps is 94% after the five first divisions. We designed a particle filter with a cutoff size of 15 μm to prevent dust particles or debris from clogging the chip. The microfluidic chips were fabricated with PDMS using standard photo- and soft-lithography methods (PDMS, Sylgard 184, Dow Chemical, USA, see supplementary text for detailed protocols). We connected the chip using PTFE tubing (1mm outer diameter), and we used a peristaltic pump to ensure media replenishment (Ismatec, Switzerland). We used standard rich media supplemented with 2% dextrose (YPD). See supplementary methods for additional details.

### Microscopy

The microscope was built from a modular microscope system with a motorized stage (ASI, USA, see the supplementary text for the detailed list of components), a 20x objective 0.45 (Nikon, Japan) lens, and an sCMOS camera (ORCA flash 4.0, Hamamatsu, Japan). A dual-band filter (#59022, Chroma Technology, Germany) coupled with a two-channel LED system (DC4104 and LED4D067, Thorlabs, USA). The sample temperature was maintained at 30°C thanks to a heating system based on an Indium Thin Oxide coated glass and an infrared sensor coupled to an Arduino-based regulatory loop. Micromanager v2.0(Edelstein et al. 2014) was used to drive all hardware, including the camera, the light sources, and the stage and objective motors. We developed a custom autofocusing routine to minimize the autofocus time (https://github.com/TAspert/DetecDiv_Data). The interval between two frames for all the experiments was 5 min. We could image approximately 80 fields of view (0.65mmx0.65mm) in brightfield and fluorescence (using a dual-band GFP-mCherry filter) with this interval. In the H_2_O_2_ stress response experiments, cells were exposed abruptly to a medium containing the desired concentration (from 0.3mM to 0.8mM) and fluorescence was acquired every 15minutes.

### Additional datasets for the comparative study of division detection

Time-lapse image datasets of individual mother cells trapped in microfluidic devices were obtained from the Murat Acar and Peter Swain lab. The datasets were used to compare the performance of our cell division tracking pipeline, as described in the main text. Data from the Acar lab were generated on a Nikon Ti Eclipse using 60x phase-contrast imaging with a 10min-interval and a single z-stack, as previously described (Liu, Young, and Acar 2015). Data from the Swain lab were obtained using a Nikon Ti Eclipse microscope using 60x phase-contrast imaging, a 2.5min-interval (Crane et al. 2014), and 5 z-stacks were combined into a single RGB image used as input to the classifier. We also used a separate trap design from our own lab that is similar to a previously reported design (Jo et al. 2015) which was imaged on a Nikon Ti Eclipse microscope using a 60x phase-contrast objective.

### Image processing

We developed a Matlab software, DetecDiv, which provides different classification models: image classification, image sequence classification, time series classification, and pixel classification (semantic segmentation), see Supplementary Table 2. DetecDiv was developed using Matlab, and additional toolboxes (TB), such as the Computer Vision TB, the Deep-learning TB, and the Image Processing TB. A graphical user interface was designed to facilitate the generation of the training sets. The DetecDiv software is available for download on GitHub: https://github.com/gcharvin/DetecDiv

#### Image classification for division tracking and lifespan reconstruction

DetecDiv was used to classify images into six classes after supervised training using a GoogleNet (Szegedy et al. 2015) network combined with an LSTM network (Hochreiter and Schmidhuber 1997). See supplementary text for details.

#### Image segmentation from brightfield and fluorescent images

DetecDiv was used to segment images using a pixel classification model called Deeplab v3+ (Chen et al. 2018), after supervised training based on 1400 and 3000 manually annotated images (respectively). See supplementary text for details.

#### Senescence Entry Point detection

DetecDiv was used to detect cell-cycle slowdown (Senescence Entry Point) from a temporal sequence of classes obtained using the division tracking network. The training was based on manual annotation of 200 lifespans. See supplementary text for details.

#### Statistics

All experiments have been replicated at least twice. Data are presented in Results and Figures as the mean ± SEM (curves) or median. Group means were compared using the Two-sample t-test. A P-value of < 0.05 was considered significant.

#### Computing time

Image processing was performed on a computing server with 8 Intel Xeon E5-2620 processors and 8 co-processing GPU units (Nvidia Tesla K80), each of them with 12Go RAM. Under these conditions, the classification of the time-series of 1000 frames from a single trap (roughly 60×60pixels) took 3.5s to the CNN/LSTM classifier. For the image segmentation, the DeepLab v3+ network took about 20s to classify 1000 images.

## Supporting information

Strain Table

## Acknowledgments

We thank Audrey Matifas for constant technical support throughout this work, Sophie Quintin and Nacho Molina for carefully reading the manuscript. We are very grateful to Murat Acar and David Moreno Fortuño, as well as Peter Swain and Julian Pietsch for providing the additional time-lapse datasets analyzed in this study. We thank Olivier Tassy for their insightful discussions. We thank Denis Fumagalli at the IGBMC Mediaprep facility for media preparation. We are grateful to the IT service for efficient support and providing the computing resources. We thank the Charvin lab members, Bertrand Vernay, Jerome Mutterer, Serge Taubert, and the IGBMC imaging facility for discussions and technical support. This work was supported by the Agence Nationale pour la Recherche (T.A. and G.C.), the grant ANR-10-LABX-0030-INRT, a French State fund managed by the Agence Nationale de la Recherche under the frame program Investissements d’Avenir ANR-10-IDEX-0002-02.

## Data Availability

Annotated datasets and trained classifiers used in this study are available for download as indicated:

- Lifespan analyses:
  - Training & test datasets: doi.org/10.5281/zenodo.6078462
  - Trained Network (CNN+LSTM): doi.org/10.5281/zenodo.5553862
- Brightfield image segmentation:
  - Training & test datasets: doi.org/10.5281/zenodo.6077125
  - Trained Network (Encoder-Decoder Deeplabv3+): doi.org/10.5281/zenodo.5553851
- Cell-cycle slowdown (Senescence Entry Point) detection:
  - Training & test datasets: doi.org/10.5281/zenodo.6075691
  - Trained Network (LSTM): doi.org/10.5281/zenodo.5553829

Information regarding the design of the microfluidic device and of the custom imaging system are available on https://github.com/TAspert/DetecDiv_Data.

## Code availability

The custom MATLAB software DetecDiv, used to analyze imaging data with deep-learning algorithms, is available on https://github.com/gcharvin/DetecDiv.

This software distribution features a tutorial on how to use the graphical user interface: https://github.com/gcharvin/DetecDiv/blob/master/Tutorial/GUI_tutorial.md

It also provides the main commands to use the DetecDiv pipeline in custom user-defined scripts:

https://github.com/gcharvin/DetecDiv/blob/master/Tutorial/commandline_tutorial.md

A demo project that contains all the necessary files to learn how to use DetecDiv can be downloaded from zenodo:

https://doi.org/10.5281/zenodo.5771536

