## Supplementary text for "DetecDiv, a generalist deep-learning platform for automated cell division tracking and survival analysis"

**Théo Aspert[1234*], Didier Hentsch[1234] and Gilles Charvin[1234*]**

*1) Department of Developmental Biology and Stem Cells, Institut de Génétique et de Biologie Moléculaire et Cellulaire, Illkirch, France*

*2) Centre National de la Recherche Scientifique, UMR7104, Illkirch, France*

*3) Institut National de la Santé et de la Recherche Médicale, U964, Illkirch, France*

*4) Université de Strasbourg, Illkirch, France*

### Supplementary Methods

##### Cell culture

For each experiment, freshly thawed cells were grown overnight, diluted in the morning, and allowed to perform several divisions (~5 hours at 30°C) before injection into the microfluidic device. Yeast extract Peptone Dextrose (YPD) medium was used throughout the experiments.

##### Microfluidic mold fabrication

The designs were created on AutoCAD (see https://github.com/TAspert/DetecDiv_Data to download the design) to produce chrome photomasks (jd-photodata, UK). Then, the microfluidic master molds were made using two rounds of classical photolithography steps.

The array of 2000 traps was created from a 5.25µm deposit by spinning (WS650 spin coater, Laurell, France) 3mL of SU8-2005 negative photoresist at 2500rpm for 30sec on a 3” wafer (Neyco, FRANCE). Then, a soft bake of 3min at 95°C on heating plates (VWR) was performed, followed by exposure to 365nm UVs at 120 mJ/cm² with a mask aligner (UV-KUB3, Kloé, FRANCE). Finally, a post-exposure bake identical to the soft bake was performed before development using SU-8 developer (Microchem, USA).

The second layer with channel motifs was made of a 30µm deposit of SU8-2025, by spinning it at 2500rpm for 30s. Subsequently, a soft bake of 3min at 65°C and 6min at 95°C was performed. The wafer was then aligned with the mask containing the motif of the second layer before a 120mJ/cm² exposure. A post-exposure bake similar to the soft bake was then performed.

After each layer, we performed a hard bake at 150°C for 15min to anneal potential cracks and stabilize the photoresist. Finally, the master molds were treated with chlorotrimethylsilane to passivate the surface.

**Microfluidics chip design, fabrication, and handling**

The microfluidic device is composed of an array of 2048 microstructures able to trap a mother cell while removing successive daughter cells, similarly to previously designed (Ryley and Pereira-Smith 2006; Zhang et al. 2012; Lee et al. 2012), (Crane et al. 2014; Jo et al. 2015; Liu, Young, and Acar 2015). The traps are composed of two symmetrical structures separated by 3µm (see Figure 1 and Figure 1 - figure supplement 1D-G), in such a way that only one cell can be trapped and remain in between the structures. We have measured that 94% of the cells that underwent at least 5 divisions in the trap would stay inside until their death. This is striking contrast with the results obtained with a device with a semi-open trap geometry ( see Figure 3 and FIgure 3 - figure supplement 3 for details). Moreover, a particle filter with a cutoff size of 15µm is present before each array of traps, preventing dust particles or debris from clogging the chip (Figure 1 - figure supplement 1C).

The microfluidic devices were fabricated using soft-lithography by pouring polydimethylsiloxane (PDMS, Sylgard 184, Dow Chemical, USA) with its curing agent (10:1 mixing ratio) on the different molds. The chips were punched with a 1mm biopsy tool (KAI, Japan) and covalently bound to a 24 × 50 mm coverslip using plasma surface activation (Diener Zepto, Germany). The assembled chips were baked for 30 min at 60°C to consolidate covalent bonds between glass and PDMS. The chip was then plugged using a 1mm Outer Diameter (O.D.) PTFE tubing (Adtech, UK) and the channels were primed using culture media for 5min. After that, cells were injected through the outlet using a 5mL syringe and a 26G needle for approximately 1 minute per channel by applying very gentle pressure. The cell filter placed upstream of the trapping area prevented the cells from entering the tubing connected to the inlet (Figure 1 - figure supplement E). Then, the inlet of each microfluidic channel was connected to a peristaltic pump (Ismatec, Switzerland) with a 5µL/min rate to ensure a constant replenishment of the media and dissection of the daughter cells (Figure 1 - figure supplement 1E). This procedure avoids potential contamination by cells forming colonies upstream of the trapping area, which would induce the clogging of the device after 1-2 days of experiment, therefore making long-lasting experiments more robust.

**Microscopy**

Microscope

The microscope was built from a modular microscope system (RAMM, ASI, USA) with trans- (Oly-Trans-Illum, ASI, USA) and epi- (Mim-Excite-Cond20N-K, ASI, USA) illumination. This microscope frame provides a cost-effective solution to build a minimal microscopy apparatus to perform robust image acquisition over several days (Figure 1 - figure supplement 1B).

It is equipped with a motorized XY stage (S551-2201B, ASI, USA), a stage controller (MS200, ASI, USA), and a stepper motor to drive the 20x N.A. 0.45 Plan Fluor objective (Nikon, Japan) and an sCMOS camera (ORCA flash 4.0, Hamamatsu, Japan) with 2048 x 2048 pixels (*i.e.*, 650 microns x 650 microns field of view at 20x magnification). We used a dual-band filter (#59022, Chroma Technology, Germany) coupled with two-channel LED illumination (DC4104 and LED4D067, Thorlabs, USA), which allows fast imaging of GFP and mCherry without any filter switching.

Sample holder and temperature control

We designed a custom 3D-printed sample holder (by extruding PLA material with a MK3S+ printer, Prusa Research, Czech republic) for the microfluidic device to ensure the mechanical stability of the microfluidic device.

In addition, we developed a custom temperature control system to maintain a constant temperature (30°C) and guarantee optimal cell growth throughout the experiment. Briefly, we used an Indium Tin Oxide (ITO) Coated glass, an electrically conductive and transparent material, in direct contact with the PDMS chip. Therefore, applying a voltage to the ITO glass allowed the Joule effect to heat the glass and the adjacent PDMS chip (Figure 1 - figure supplement 1A).

To achieve a temperature control loop, we used an infrared sensor attached to the objective and facing towards the bottom glass coverslip in contact with the cells. The sensor allowed in situ temperature measurement and was used in an Arduino-based PID control loop to regulate the heating power to maintain the setpoint temperature. About 0.3W was sufficient to maintain a 30°C temperature at room temperature. Notably, the temperature profile obtained with this method was homogenous and constant throughout the experiment. Furthermore, the glass is fully transparent to visible light. In addition, it does not interfere with fluorescent light when using an inverted microscope since it is located at the end of the optical path, after the sample.

Software and time-lapse acquisition parameters

Micromanager v2.0. was used to drive the camera, the light source, the XYZ controller, and the LED light source for fluorescence epi-illumination. We developed a specific program in order to drive the temperature controller from the Arduino (The source code is available on github: https://github.com/TAspert/ITO_heating_device).

Unless specified otherwise, the interval between two brightfield frames for all the experiments was 5minutes, and images were recorded over 1000 frames (*i.e.*, ~3 days). We used three z-stack for brightfield imaging (spaced by 1.35 microns) to ease the detection of small buds during the image classification process for cell state determination. Fluorescent images were acquired with a 10min interval using 470nm illumination for 50ms. Up to 80 fields of view were recorded over the 5min interval.

Autofocusing

To keep a stable focus through the whole experiment, we developed a custom software-based autofocus routine that finds the sharpest image on the first field of view and then applies the focus correction to the rest of the positions. This method provides faster scanning of all fields of view in a reasonable time. Nevertheless, it is almost as efficient as performing autofocusing on each position since the primary source of defocusing in our setup is the thermal drift, which applies identically to all the positions.

#### Image processing

DetecDiv software

We designed a custom software, DetecDiv, as a Matlab backend to provide an expandable interface to perform microscopy data processing, based on existing resources in Matlab's Deep Learning Toolbox and Computer Vision Toolbox. DetecDiv can be used with an arbitrarily large number of classes, image channels, types, and sizes (Table 6). Therefore, its capabilities go far beyond the scope of this study.

First, DetecDiv offers graphical tools to identify regions of interest (referred to as ROIs throughout the manuscript) in image sequences. In addition, a cross-correlation function can be used to automatically detect similar regions in images, such as the traps in the microfluidic device (Figure 1).

Then, several classification models can be defined to process the images: 1) Image classification using a convolutional network (CNN, see Figure 2 - figure supplement 1); 2) a combined CNN/LSTM classifier (Figure 2A); 3) an LSTM network to perform sequence-to-sequence (as in Figure 4A) or sequence-to-one classification; 4) An encoder/decoder classifier to perform pixel classification (semantic segmentation) based on the Deeplab v3+ architecture (Chen et al. 2018), (Figure 5); 5) Similar routines as in 1-4, but for regression analyses (not used in the present study, see Supplementary Table 6 for the detailed list of available networks). DetecDiv allows the user to choose among several CNNs -such as GoogleNet or Resnet50 for all image classifications/segmentations applications.

DetecDiv provides a graphical user interface to generate the ground truth required for both training and testing the classifiers used in the image classification, pixel classification, and time-series classification pipelines. Furthermore, we paid attention to making this step as user-friendly as possible. For instance, we used keyboard shortcuts to assign labels to individual frames (it takes about 5-10 min to annotate 1000 frames in the case of cell state assignment). Similarly, direct "painting" of objects with a mouse or a graph pad can be used to label images before launching the training procedure for pixel classification.

DetecDiv training and validation procedures are run either at the command line, which allows using remote computing resources, such as a CPU/GPU cluster, or using a Matlab GUI application. All the relevant training parameters can be easily defined by the user. We designed generic routines to benchmark the trained classifiers that allow an in-depth evaluation of the classifiers' performances (Laine et al. 2021). Trained classifiers can be exported to user-defined repositories and classified data can be further processed using custom Matlab scripts, and images sequences can be exported as .mat or .avi video files.

Last, DetecDiv provides additional post-processing routines to extract generations and lifespan data for further analysis, as performed in the present study.

Convolutional Neural Networks (CNN) for classification of the cellular budding status and death

We used an image classifier to assess the state of cells in the cell cycle (small, large-budded, etc.) using brightfield images of individual traps. For each frame, we combined the three z-stack images described above into a single RGB image, which was used as input for the classifier.

We defined six classes, four of which represent the state of the cell, i.e., unbudded (‘unbud’), small-budded (‘small’), large-budded (‘large’), dead (‘dead’), as shown in Figure 2. Two additional classes are related to the state of the trap: trap with no cell (‘empty’) and clogged trap (‘clog’).

The small class was attributed to images on which the mother cell displayed a bud below a certain threshold size. This size was defined to represent approximately half of the cell-cycle images, but also to stay below the smallest daughter cells.

The large class was attributed to images on which the mother cell displayed a bud above the aforementioned threshold size. If no bud was visible at the mother surface after the cytokinesis, the image was labeled as "large" if the daughter cell remained in contact with the mother, or "unbudded" if the mother cell was left alone in the trap.

An image was labeled as ‘dead’ when the mother cells appeared as dead. This includes an unambiguous, very abrupt (within one frame) and strong change in contrast and appearance of the cell.

The empty class was attributed to images on which no cell was present in the ~ three first quarters (from the bottom) part of the trap.

Finally, the clogged class was used for images where more than ~ 50% of the outer space of the trap was filled with cells.

We trained a pre-trained Inception convolutional neural network (Szegedy et al. 2015) to classify images according to these six classes using a training set of 200000 representative manually annotated brightfield images (*i.e.*, 200 traps monitored during 1000 frames). The training of the classifier was achieved using Adaptive Moment estimation (Adam) optimizer (Kingma and Ba 2015). Specific parameters used can be found in the supplementary table T2.

After the training procedure, we tested the classifier using a dataset composed of 50 independent cell traps (*i.e.* ~50000 images) that were manually annotated and used for benchmarking (Figure 2 - figure supplement 2).

<u>Cell-cycle duration measurements and replicative lifespan (RLS) reconstruction based on classification results</u>

As the image classifier outputs a label for each frame corresponding to one of the 6 classes defined above, we used the sequence of labels to reveal the successive generations of the cells: the oscillations between the “large” and “small” or “unbudded” and “small” classes captured the entry into a new cell cycle (i.e. a budding event).

The first occurrence of one of the four following rules was used as a condition to stop the lifespan: 1) the occurrence of a “Dead” class; 2) division arrest for more than 10 hours; 3) occurrence of a “Clogged” class; 4) the occurrence of an “Empty” class (the mother left the trap Figure 1 - figure supplement 1G and Figure 3 - figure supplement 1), which was a rare case. Premature lifespan arrests due to clogging or mother cell removal were not considered further for lifespan analyses.

This set of rules was used to compute the cell-cycle duration and the RLS of each individual cell when using either the CNN or the combined CNN/LSTM architecture (see below). However, in order to improve the accuracy of the method based only on the CNN, we implemented an additional “post-processing” step (referred to as PP in Figure 2), namely that two consecutive frames with a “dead” label are necessary to consider a cell as dead.

Image sequence classification using combined CNN and a long short-term memory network (LSTM)

To provide a more accurate classification of the image according to the cellular state, we added a bidirectional long short-term memory (LSTM) network with 150 hidden units to the CNN network (Hochreiter and Schmidhuber 1997). The LSTM network takes the whole sequence of images as input (instead of independent images in the case of the CNN). 200 ROIs (with 1000 frames each) were used to train the LSTM network independently of the CNN network (see training parameters in Supplementary table T3 and benchmarks on Figure Figure 2 - figure supplement 2B-D). The CNN and the LSTM network were then assembled as described in Figure 2A in order to output a sequence of labels for each sequence of images. The assembled network was then benchmarked using a set of 50 independent annotated ROIs, as described above.

The training & test datasets are available at: doi.org/10.5281/zenodo.6078462

The trained network is available at: doi.org/10.5281/zenodo.5553862

Assessment of cell-cycle slowdown using an LSTM network

We designed a time series classification method to identify when the cell cycle starts to slow down (Senescence Entry Point, or SEP, see main text). For this, we trained a bidirectional LSTM network with 150 hidden units to classify all the frames in each lifespan, into two classes, ‘pre-SEP and ‘post-SEP’, using a manually annotated dataset containing 200 ROIs. To achieve these annotations, we designed a custom annotation GUI allowing us to monitor the successive states of a mother cell of interest over time, as output by the CNN/LSTM network above. Thi tool was convenient to detect the cell cycle slow down occurring upon entry into senescence. Then, the LSTM network was trained using class probabilities from the previously described CNN/LSTM (unbudded, small, large, dead), see Figure 4 - figure supplement 1 for benchmarking results on a test set with 47 ROIs.

The training and test datasets are available at: doi.org/10.5281/zenodo.6075691

The trained network is available at: doi.org/10.5281/zenodo.5553829

Brightfield and fluorescence images semantic segmentation using DeepLab v3+

Cell and nuclei contours were determined based on brightfield and fluorescence images, respectively, using the deep learning-based semantic segmentation architecture DeepLab v3+ (Chen et al. 2018). We used 3 classes (*i.e.*, background, mother cell, other cells) for cell segmentation to distinguish between the mother cell and its surrounding buds or daughter cells, see Figure 4 and S7). For nuclei segmentation, only two classes were defined (*i.e.*, background and nucleus, see Figure 5 and Figure 5 - figure supplement 1). We used the Resnet50 CNN in the encoder part of the network because it provided better segmentation results than Googlenet. We trained the network using a manually curated training set of 1400 and 3000 images for brightfield and fluorescence images, respectively (see Figure S7 and S8 for benchmarking results). Specific parameters used can be found in the supplementary table T4 (cell segmentation) and T5 (nucleus segmentation). In addition, we have implemented a cross-validation routine to test the sensitivity of the classifier used for cell segmentation to the training and test datasets. For this purpose, we have performed 30 successive random draws of 200 annotated ROIs to be used as a training set and 50 annotated ROIs for testing the classifier (the total number of manually annotated ROIs is 250). For each draw, we have measured the performance of the classifier (*i.e.* accuracy, recall, $F_1$-score), see Figure 5 - figure supplement 2E.

The training & test datasets are available at: doi.org/10.5281/zenodo.6077125

The trained network is available at: doi.org/10.5281/zenodo.5553851

Classifier benchmarking

We used standard benchmarking to estimate the efficiency of image and pixel classifiers. For each classifier, we computed the confusion matrix obtained by comparing the “ground truth” of manually annotated images (or time series) taken from a test set unseen by the network during training to the predictions made by the classifier. We computed the accuracy, recall, and $F_1$-score for each class. In the specific case of pixel classification (semantic segmentation), we computed these benchmarks for different values of prediction thresholds used to assign the “mother” and “nucleus” classes, as reported in Figure 5 - figure supplement 2 and 3. Then, we performed the segmentation of images using the threshold value that maximizes the $F_1$-score (0.9 and 0.35 for brightfield and fluorescence image classification, respectively).

To benchmark the detection of new generations, we used a custom pairing algorithm to detect false positive and false negative new generation events. Using this, we could compute the accuracy and recall of the detection of new generation events, and plot the correlation between paired new generations events (Figure 2D).

Statistics

All experiments have been replicated at least twice. Error bars represent the standard error-on-mean unless specified otherwise. Results of specific statistical tests are indicated in the figure legends.

#### Supplementary figures legends

**Figure 1**

**Figure supplement 1: Experimental setup and microfluidic device**

A) Schematics of the custom imaging setup built for DetecDiv (see supplementary text for details).
B) Picture of the imaging setup
C) Schematics of the microfluidic device with 16 independent channels. Each channel has one inlet, a dust filter, and one outlet.
D) Schematics of the array of cell traps. Dimensions are in microns. Inset represents close-ups on indicated areas.

E) Principle of the cell barrier used to prevent the cells from moving towards the inlet when loading the cells from the outlet. Any cell upstream of the cell array may lead to the formation of colonies hence clog the device over time.
F) Principle of the automated dissection of daughter cells: the mother is retained within the trap but their successive daughters are flushed away due to constant medium flow. Daughters may either exit the trap from the top of the bottom opening.
G) Unlike previous cell trap geometries ("classical"), the current design ("new") features shallow PDMS walls that can be deformed by large cells, hence ensuring the long-term retention of the cells. Two small claws on each side of the trap entrance further enhance retention. The retention, measured as the number of cells staying inside the trap before their death (or more than 5000min, *i.e.* the duration of the experiment), is displayed for both type of traps (N=100).

**Figure 2**

**Figure supplement 1: Principles of division tracking and lifespan reconstruction using a CNN-based image classification**

A) In this framework, the sequence of images is processed by a GoogleNet CNN that processes each image separately. The CNN extracts image features that are used to assign a label to each image among six possible classes (see supplementary methods for details). As with the CNN/LSTM architecture described in Figure 2, the sequence of labels is used to assign new generation events and the occurrence of cell death.
B) Typical sequence of label (Top) and the associated generations extracted from it (Bottom), from the image sequence of a ROI. The groundtruth is depicted in black and grey and the output of the CNN-based image classification is shown in purple. The yellow arrows indicate prediction errors. A large-to-small error leads to a false positive new generation, while a false death leads to a precocious shortening the sequence of generations.

**Figure supplement 2: Class definition and image classification benchmarks**

A) Sample images indicating how each class was defined. "Empty": empty trap, i.e., no cell; "Unbud": unbudded cell; "Small": small-budded cell; "Large": large-budded cell; "Dead": dead cell; "Clogged": clogged trap.
B) Confusion matrix obtained with a test dataset (50 trapped cells followed over 1000 frames) using the CNN image classifier. Each number in the matrix represents the number of detected events.
C) Bar plot showing the recall, accuracy, and $F_1$-score metrics obtained on each class for the CNN image classifier on the test dataset.
D) Same as B), but for the combined CNN/LSTM architecture
E) Same as C), but for the combined CNN/LSTM architecture

**Figure supplement 3: Example of image classification correctly labeling the state of the mother cell, despite the presence of surrounding cells with potentially different states.**

Top: Four images of a mother cell in contact with another cell, annotated (groundtruth) as “Small”, and classified as such. The presence of this cell does not affect the classification of the budding state of the mother cell. The green arrows indicate the actual small bud from the mother cell.
Bottom: Four images of a mother cell in contact with another cell, annotated (groundtruth) as “Large”, and classified as such. The presence of this cell does not affect the classification of the budding state of the mother cell. The red arrows indicate a small bud from the neighbor cell that could have misled the classifier.

**Supplementary movie 1: Comparison of groundtruth versus classifier predictions for the CNN/LSTM classification of the cellular state**
The left column represents the class predictions made by the CNN/LSTM classifier, while the right column represents the ground truth (determined by manual annotation). The two numbers represent the number of buds generated by the cells according to the classifier predictions and manual annotation, respectively.

**Figure 3**
**Figure supplement 1:**
Top: Time-lapse images of traps from the Acar lab (Liu, Young, and Acar 2015) from which the automated division detection can be impaired due to multiple cells in the trap (orange arrows)
Bottom: Time-lapse images of cup-shaped traps (Jo et al. 2015) from which the automated RLS analysis can be impaired due to mother/daughter replacement (red arrows).

**Figure 4**
**Figure supplement 1: Classification benchmarks for the detection of the onset of Senescence Entry using an LSTM sequence-to-sequence classification**

A) Confusion matrix obtained with a test dataset (50 time-series based on the cellular state probabilities output by the CNN/LSTM classifier) using a trained LSTM classifier. Each number in the matrix represents the number of detected events.
B) Bar plot showing the recall, accuracy, and $F_1$-score metrics obtained on each class for the LSTM classifier on the test dataset.

**Figure 5**
**Figure supplement 1: Principles of the pipeline used for semantic segmentation with DeepLab V3+**
Brightfield or Fluorescence images are separately processed by the DeepLab V3+ encoder/decoder network (Chen et al. 2018) that has been modified to classify image pixels according to user-defined classes (mother /other /background & nucleus/background for brightfield and fluorescence images, respectively). A weighted classification layer is used to deal with class imbalance.

**Figure supplement 2: Benchmarks for the semantic segmentation of brightfield images**

A) Accuracy/Recall tradeoff plot obtained by varying the output prediction threshold for the class “mother” using a test dataset that contains 50 image sequences with 1000 frames. The yellow dot indicates the point that maximizes the $F_1$-score.
B) Evolution of $F_1$-score as a function of the output prediction threshold (computed on the [0.2 : 0.95] interval. A threshold value of 0.9 maximizes the $F_1$-score (90%).
C) Confusion matrix obtained with the test dataset using a 0.9 prediction threshold.
D) Bar plot showing the recall, accuracy, and $F_1$-score metrics obtained on each class for the pixel classifier on the test dataset.
E) Cross-validation of the classification model used for semantic segmentation; Class-averaged recall, accuracy, and $F_1$-score plotted as a function of the index of the draws performed, as indicated on the legend (200 ROIs and 50 ROIs are randomly selected for training and testing upon each draw, respectively).

**Figure supplement 3: Benchmarks for the semantic segmentation of fluorescence images**

A) Accuracy/Recall tradeoff plot obtained by varying the output prediction threshold for the class “nucleus” using a test dataset that contains 25 image sequences with 1000 frames. The orange dot indicates the point that maximizes the $F_1$-score.
B) Evolution of $F_1$-score as a function of the output prediction threshold (computed on the [0.2 : 0.95] interval. A threshold value of 0.35 maximizes the $F_1$-score (91%).
C) Confusion matrix obtained with the test dataset using a 0.35 prediction threshold.
D) Bar plot showing the recall, accuracy and $F_1$-score metrics obtained on each class for the pixel classifier on the test dataset.

**Supplementary movie 1: Sample movies of individual cells following cellular state classification, cell, and nuclear contour segmentation**
The left column represents the cellular state according to the prediction made by the CNN/LSTM classifier. The middle column shows the brightfield image along with mother cell contours obtained by a semantic segmentation classifier. The right column displays the Htb2-NeonGreen fluorescence channel, along with cells contours and nuclear contours obtained by a semantic segmentation classifier.

#### Supplementary table legends

**Supplementary table 1:** List of yeast strains used in the study.

**Supplementary table 2:** Parameter values used for training the CNN+LSTM classifier (relates to Figure 2).

**Supplementary table 3:** Parameter values used for training the SEP detection classifier (relates to Figure 4).

**Supplementary table 4:** Parameter values used for training the classifier dedicated to cell segmentation (relates to Figure 5).

**Supplementary table 5:** Parameter values used for training the classifier dedicated to nucleus segmentation (relates to Figure 5).

**Supplementary table 6 :** List of all available classification models available in DetecDiv
