## Supplementary Table T6 for "DetecDiv, a generalist deep-learning platform for automated cell division tracking and survival analysis"

**Supplementary table 6: Available classification models**

| Type | Description | Input | Output | Reference figures in article |
|---|---|---|---|---|
| **Models used in the present article** | | | | |
| Image classification | CNN : GoogleNet, Resnet50, etc. | Individual images | M user-defined classes | Fig. 2 ("CNN"), 2S1-2. Table T2 |
| Image sequence classification | Combined CNN/LSTM network for sequence-to-sequence classification | Sequence of N images | N (frames) x M (user-defined classes) | Fig. 2 ("CNN/LSTM"), and 2S2. Table T2 |
| Timeseries classification | LSTM network for sequence-to-sequence classification | N (frames) x L vector | N (frames) x M (user-defined classes) | Fig. 4 and 4S1. Table T3 |
| Pixel classification (semantic segmentation) | Deeplab v3+ (with GoogleNet, Resnet50, etc. encoder) | Individual images | Labeled images (M user - defined classes) | Fig. 5, 5S1-3. Table T4 and T5 |
| **Additional models available** | | | | |
| Image sequence classification | Combined CNN/LSTM network for sequence-to-one classification | Sequence of N images | M (user-defined classes) | N/A |
| Image regression | CNN : GoogleNet, Resnet50, etc. | Individual images | 1 number | N/A |
| Image sequence regression | Combined CNN/LSTM network for sequence-to-sequence regression | Sequence of N images | N (frames) x 1 number | N/A |
| Timeseries classification | LSTM network for sequence-to-one classification | N (frames) x L vector | M (user-defined classes) | N/A |
| Timeseries regression | LSTM network for sequence-to-sequence regression | N (frames) x L vector | N (frames) x 1 number | N/A |
| Timeseries regression | LSTM network for sequence-to-one regression | N (frames) x L vector | 1 number | N/A |
