## Supplementary figures and images for "DetecDiv, a generalist deep-learning platform for automated cell division tracking and survival analysis"

### Figure 1 - supplemental Figure 1

##### Figure 1 - figure supplement 1

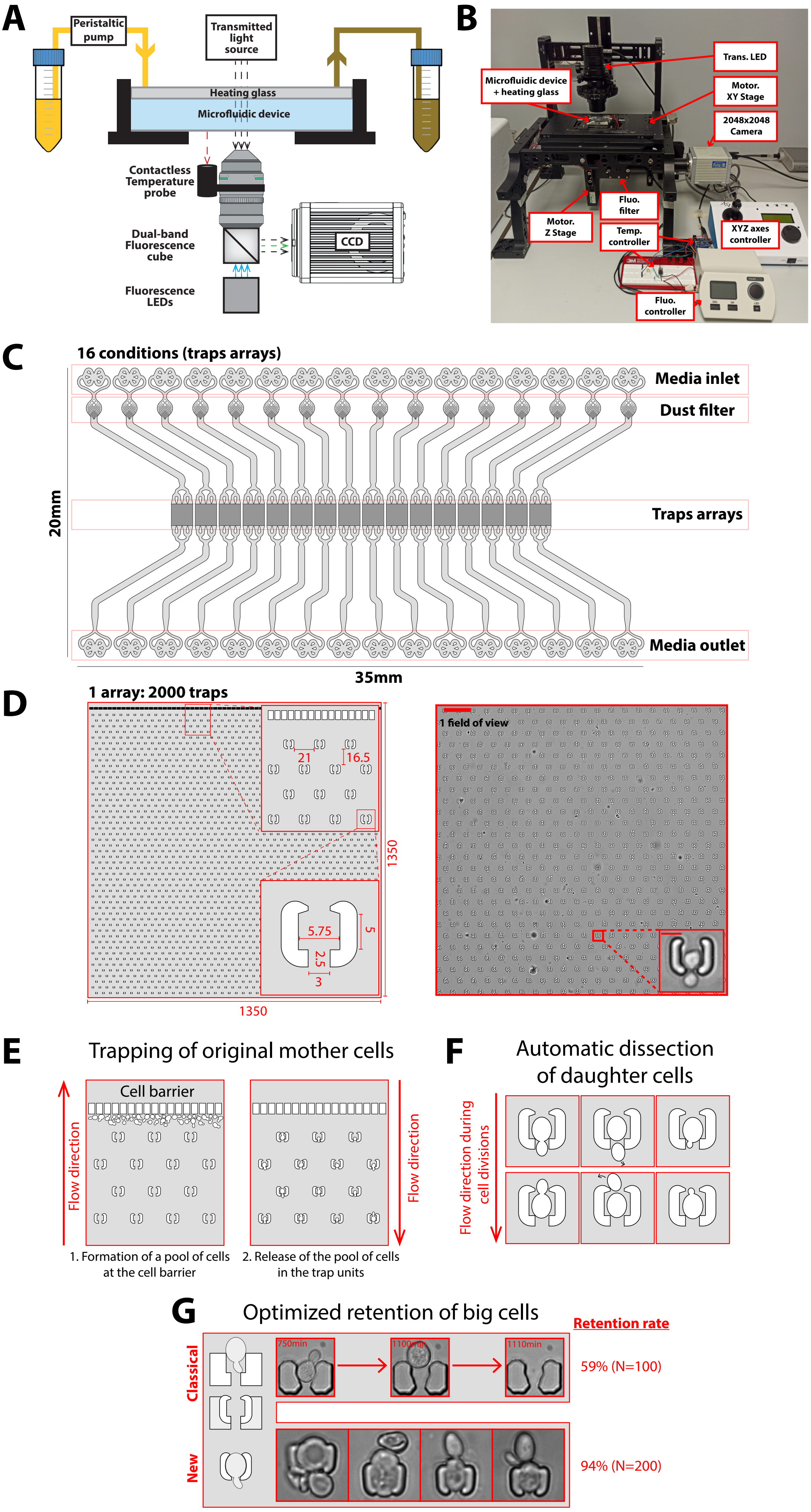

### Figure 2 - supplemental Figure 1

#### Figure 2 - figure supplement 1

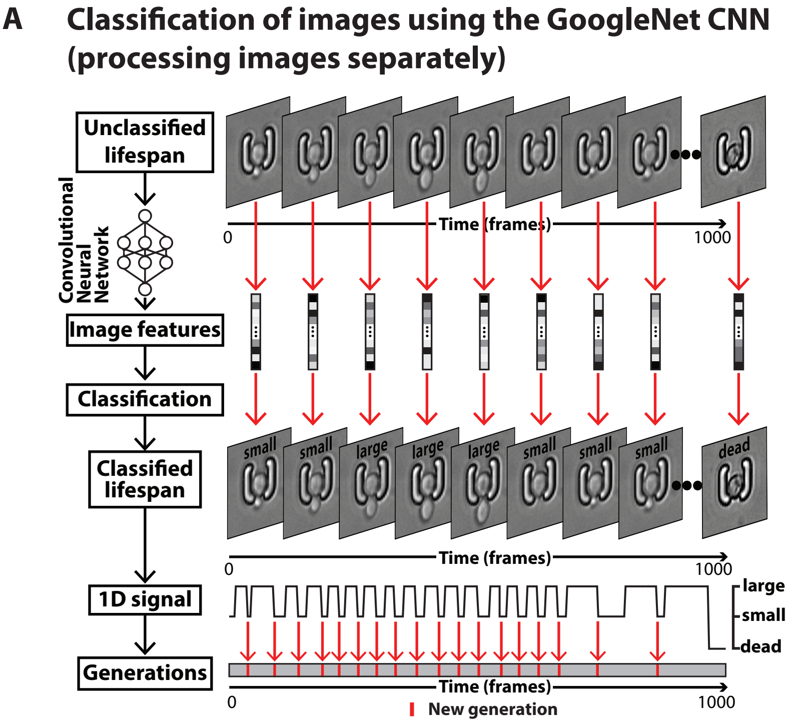


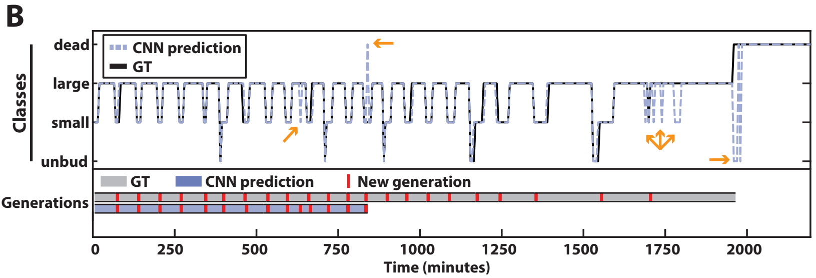

### Figure 2 - supplemental Figure 2

**Figure 2 - figure supplement 2**

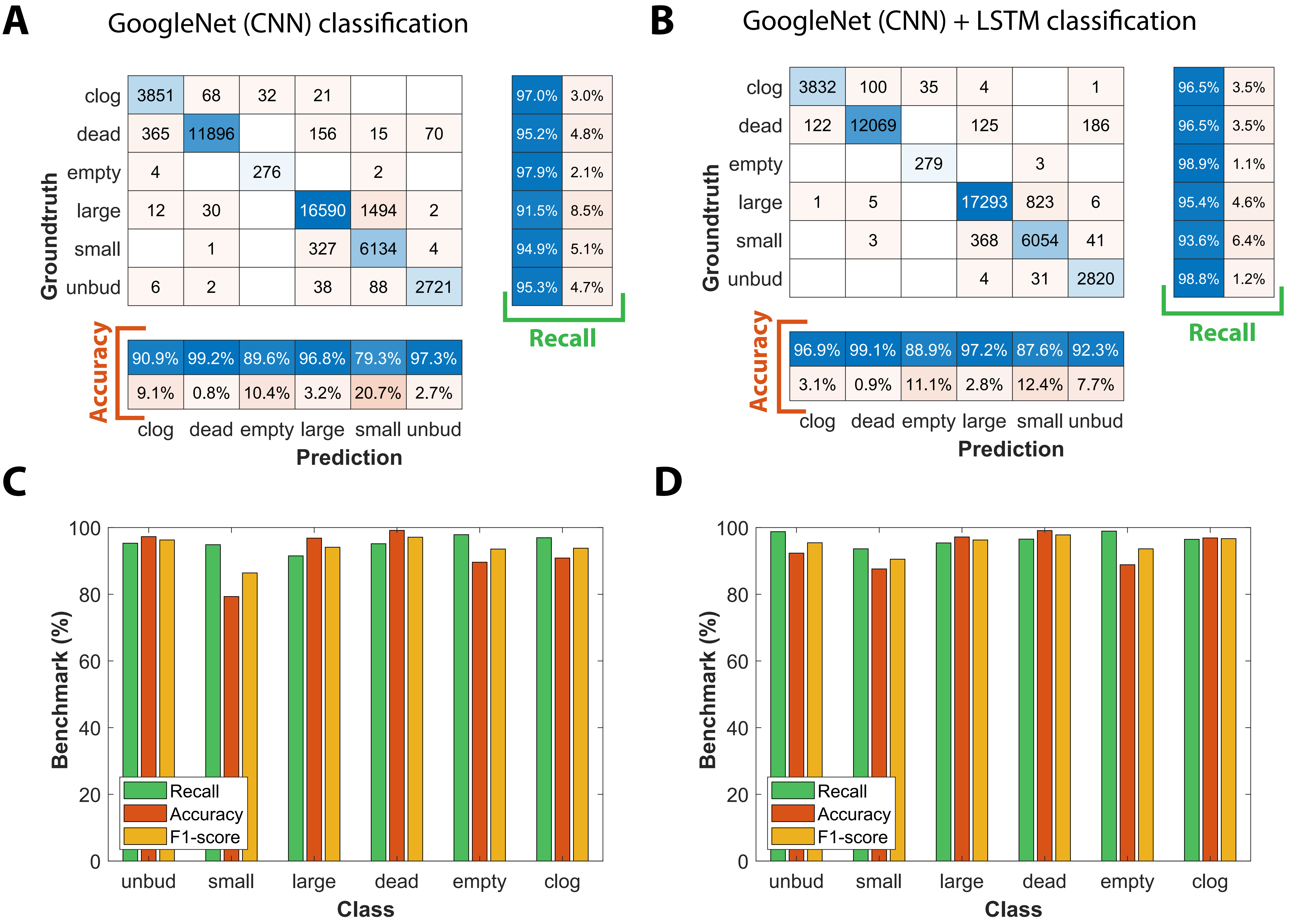

### Figure 2 - supplemental Figure 3

#### Figure 2 - figure supplement 3

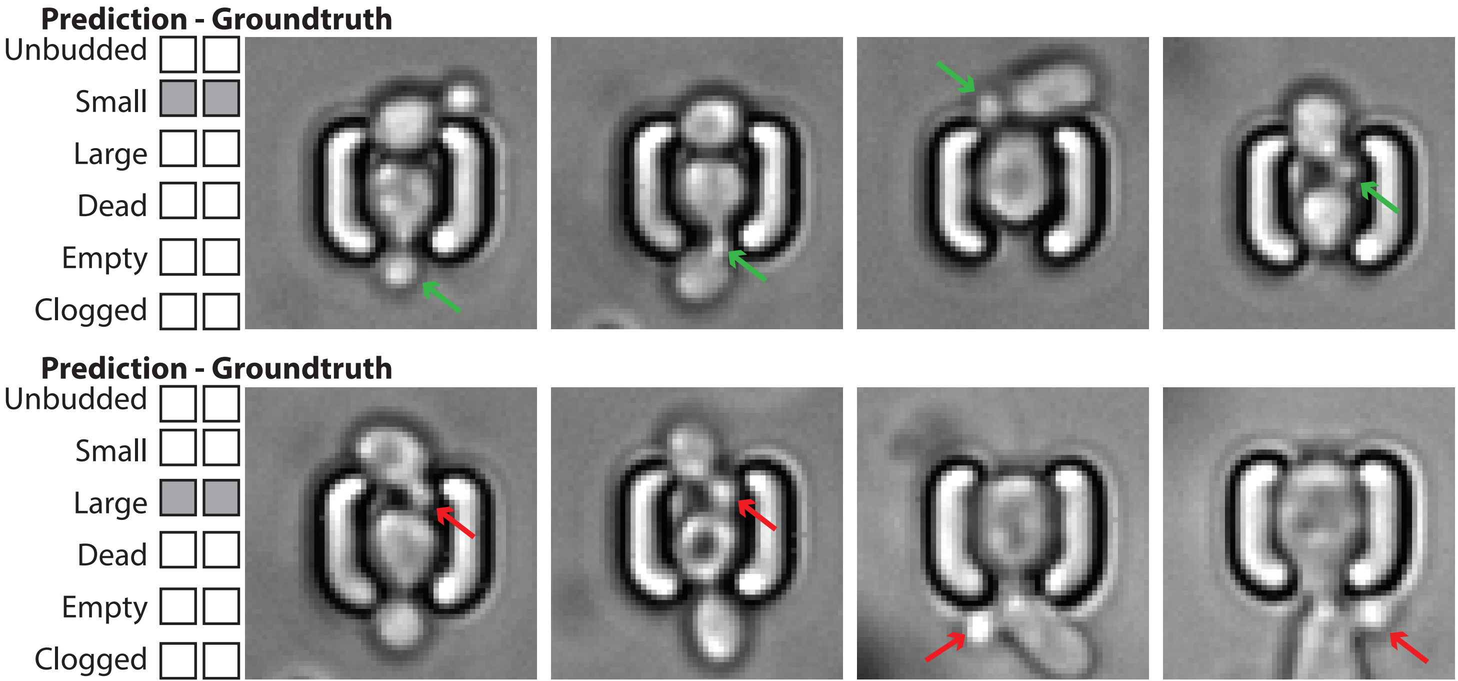

### Figure 3 - supplemental Figure 1

#### Figure 3 - figure supplement 1

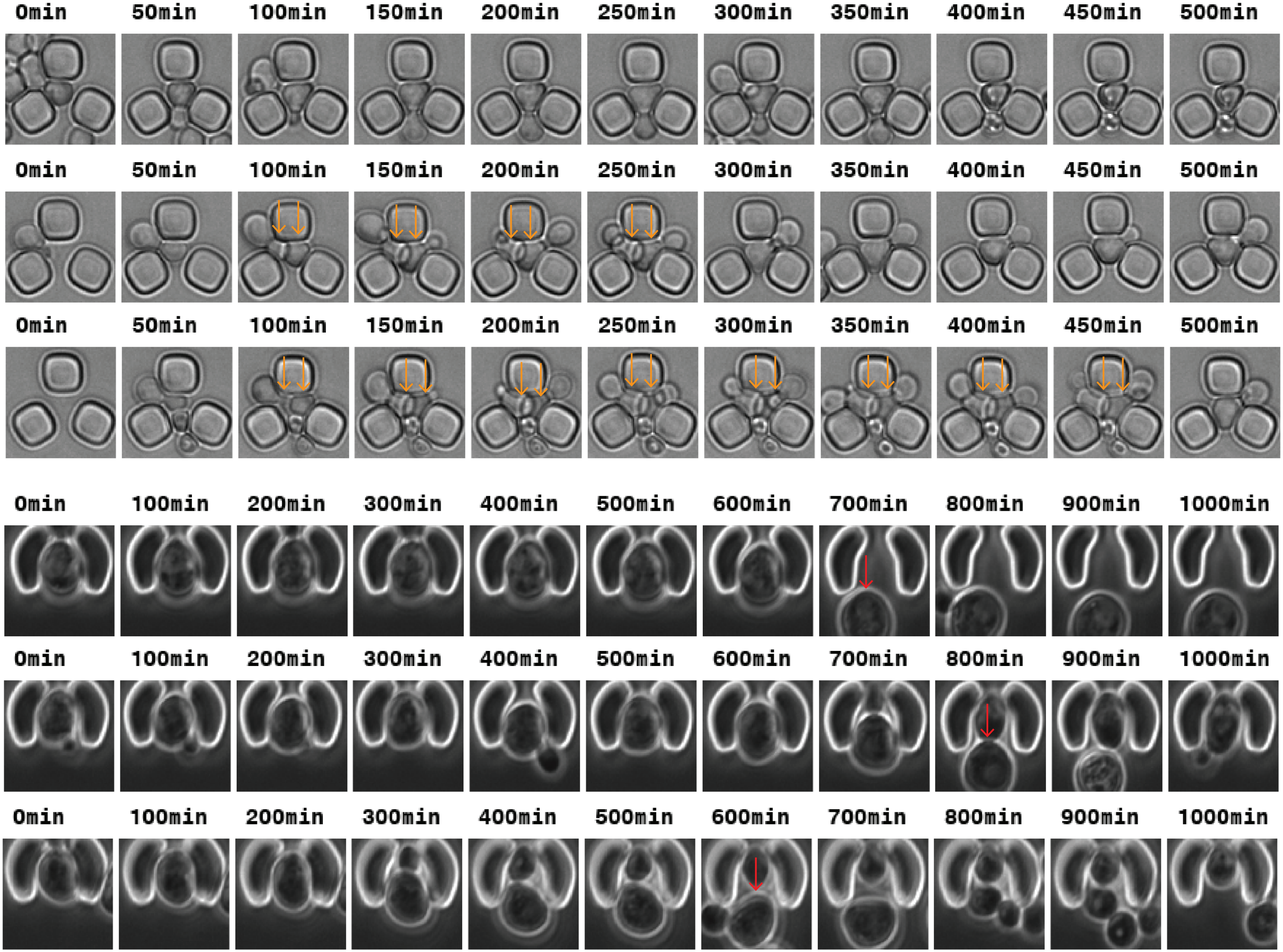

### Figure 4 - supplemental Figure 1

#### Figure 4 - figure supplement 1

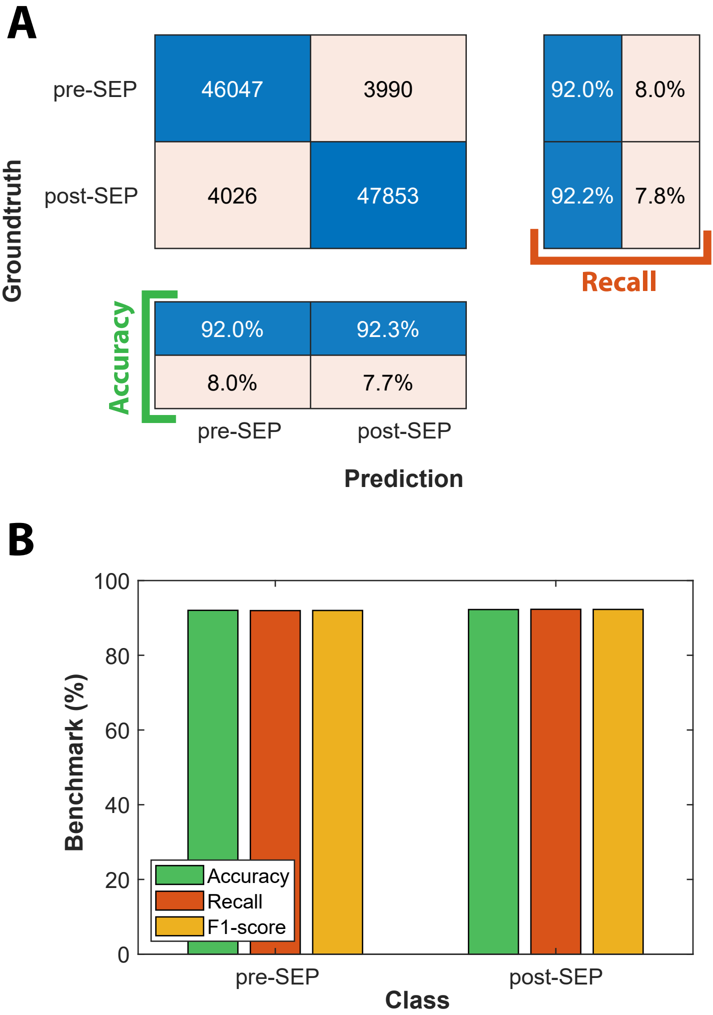

### Figure 5 - supplemental Figure 2

Figure 5 - figure supplement 2

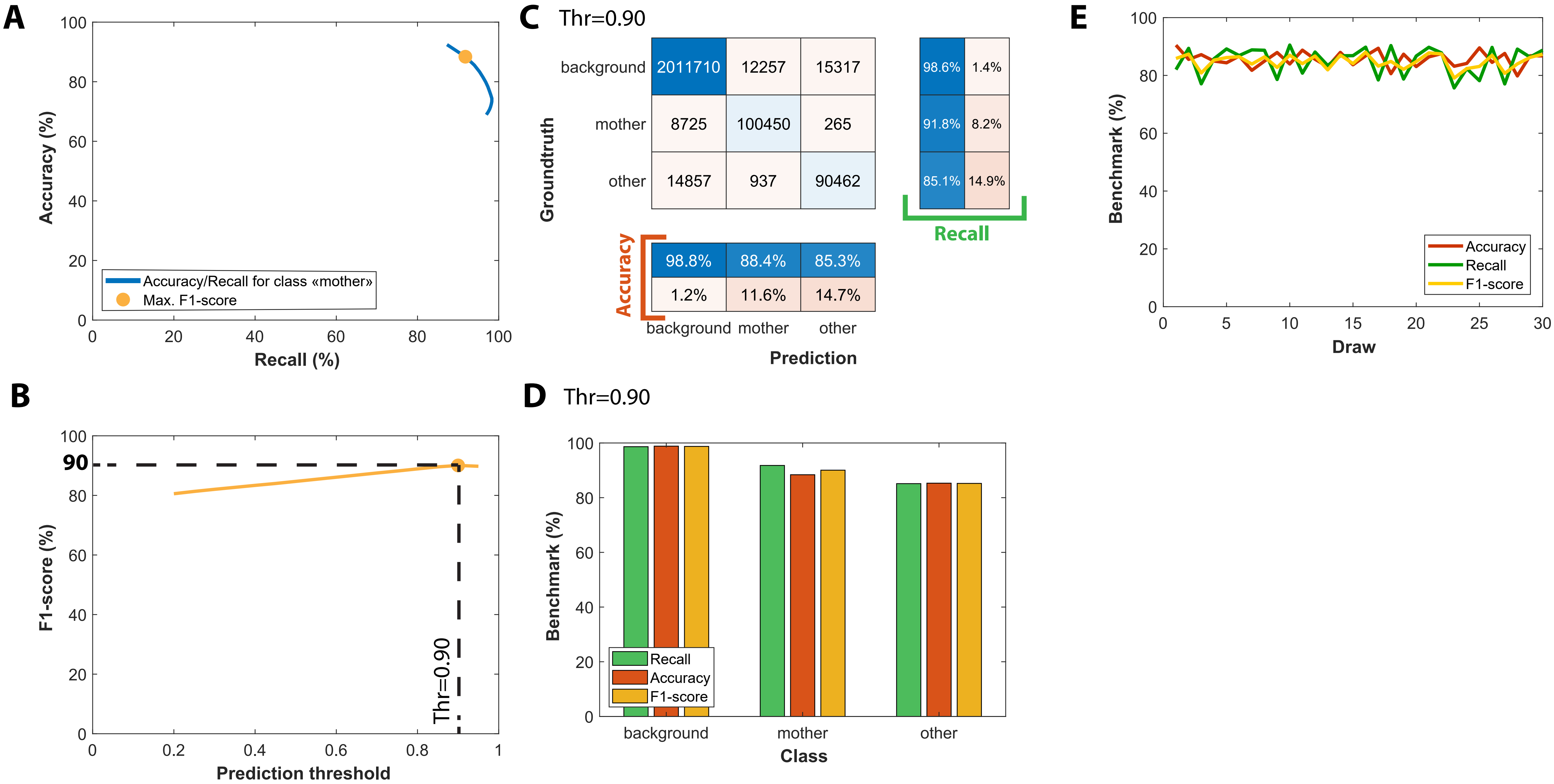

### Figure 5 - supplemental Figure 3

Figure 5 - figure supplement 3

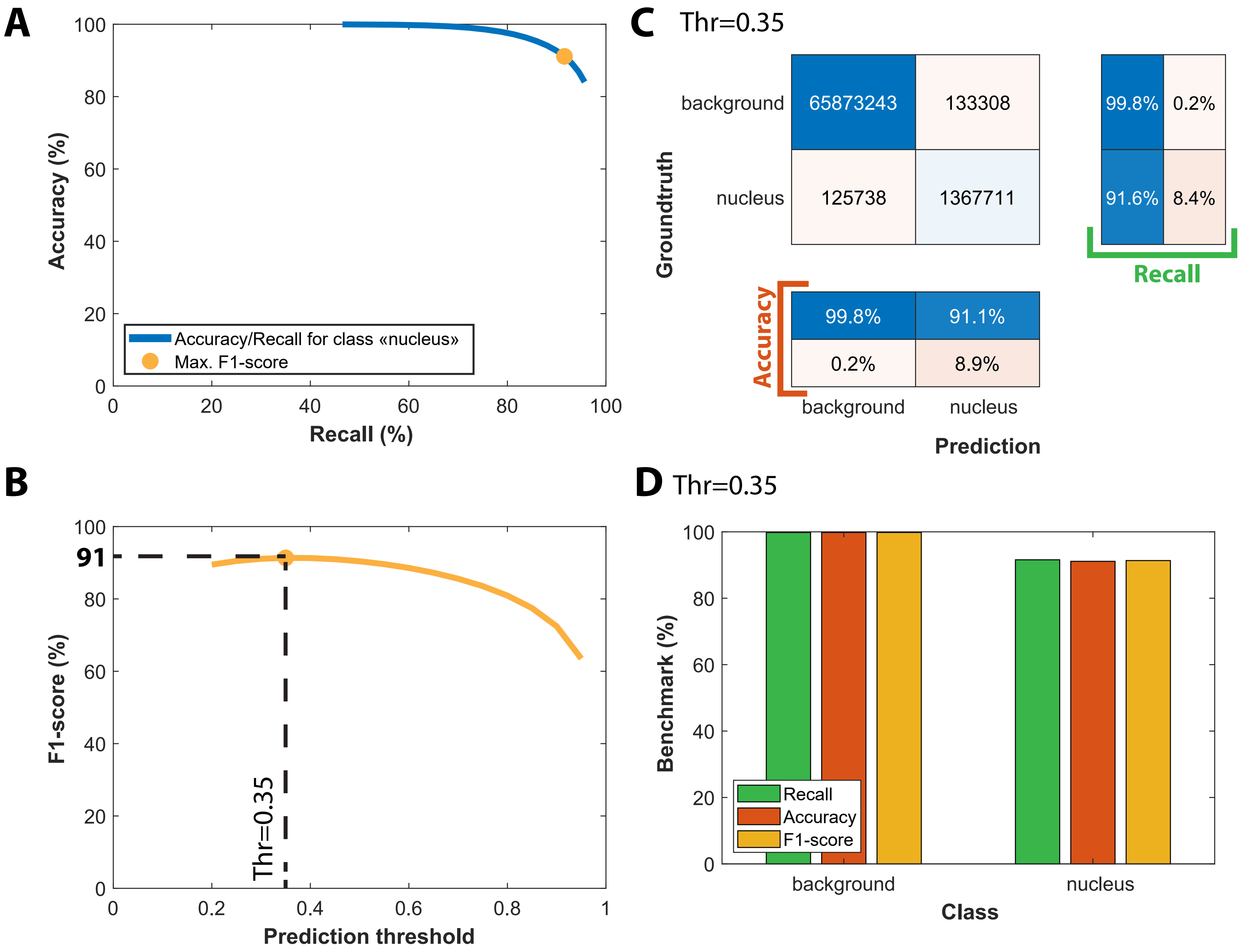
