## Supplementary material for "DetecDiv, a generalist deep-learning platform for automated cell division tracking and survival analysis": Figure 5 - supplemental Figure 1

**Figure 5 - figure supplement 1**

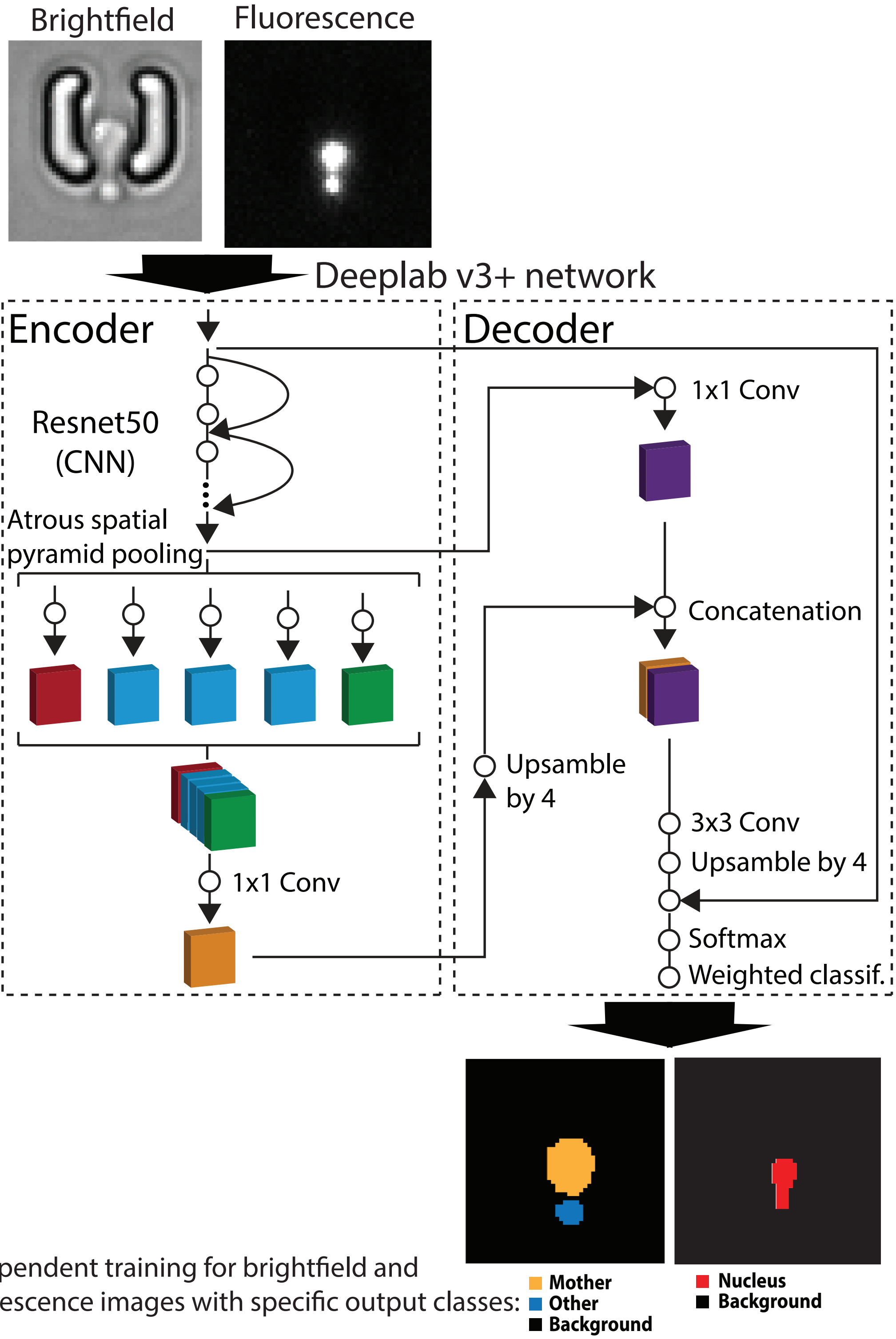


Independent training for brightfield and fluorescence images with specific output classes:
